# Toward Computationally Complete Spatial Omics

**DOI:** 10.64898/2026.03.03.709303

**Authors:** Wei Li, Liran Mao, Yunhe Liu, Fuduan Peng, Nadja Sachs, Wenrui Wu, Stephanie Pei Tung Yiu, Hanying Yan, Amelia Schroeder, Xiaokang Yu, Kaitian Jin, Shunzhou Jiang, Zihao Chen, Melanie L. Loth, Lorena Gomez, Idania Lubo, Niklas Blank, Laith Z. Samarah, Ankit Basak, Ye Won Cho, Chia-Yu Chen, David M. Kim, Alex K. Shalek, Luisa Maren Solis Soto, Joshua D. Rabinowitz, Muredach P. Reilly, Xuyu Qian, Christoph A. Thaiss, Lars Maegdefessel, Linghua Wang, Humam Kadara, Sizun Jiang, Yanxiang Deng, Mingyao Li

**Affiliations:** Statistical Center for Single-Cell and Spatial Genomics, Department of Biostatistics, Epidemiology and Informatics, Perelman School of Medicine, University of Pennsylvania, Philadelphia, PA, USA; Department of Pathology and Laboratory Medicine, Perelman School of Medicine, University of Pennsylvania, Philadelphia, PA, USA; Graduate Group in Genomics and Computational Biology, Perelman School of Medicine, University of Pennsylvania, Philadelphia, PA, USA; Department of Genomic Medicine, University of Texas MD Anderson Cancer Center, Houston, TX, USA; Department for Vascular and Endovascular Surgery, TUM Klinikum Rechts der Isar, Technical University of Munich, Munich, Germany; German Center for Cardiovascular Research (DZHK), Berlin, Germany; Partner Site Munich Heart Alliance; Center for Virology and Vaccine Research, Beth Israel Deaconess Medical Center, Harvard Medical School, Boston, MA, USA; Department of Translational Molecular Pathology, The University of Texas MD Anderson Cancer Center, Houston, TX, USA; Arc Institute, Palo Alto, CA, USA; Department of Pathology, Stanford University, Stanford, CA, USA; Department of Chemistry, Princeton University, Princeton, NJ, USA; Lewis-Sigler Institute of Integrative Genomics, Princeton University, Princeton, NJ, USA; Ludwig Institute for Cancer Research, Princeton University, Princeton, NJ, USA; Department of Chemistry, Massachusetts Institute of Technology, Cambridge, MA, USA; Department of Oral Medicine, Infection and Immunity, Harvard School of Dental Medicine, Boston, MA, USA; Division of Cardiology, Department of Medicine, Vagelos College of Physicians and Surgeons, Columbia University, New York, NY, USA; Irving Institute for Clinical and Translational Research, Columbia University Irving Medical Center, New York, NY, USA; Department of Pediatrics, Children’s Hospital of Philadelphia, University of Pennsylvania Perelman School of Medicine, Philadelphia, PA, USA; Department of Medicine, Karolinska Institutet, Stockholm, Sweden; Epigenetics Institute, Perelman School of Medicine, University of Pennsylvania, Philadelphia, PA, USA

**Keywords:** Virtual tissue model, Multimodal spatial omics, Integration, Prediction, Deep learning

## Abstract

Multimodal spatial omics has transformed biology by mapping molecular complexity within intact tissues, yet current technologies remain limited in the number of modalities measured simultaneously and often produce lower-quality data than single-modality assays. We present COSIE, a computational framework that generates high-resolution, multilayered molecular landscapes across tissue sections, individuals, and platforms. COSIE integrates histology, epigenome, transcriptome, proteome, and metabolome into a unified representation. Applied to 12 datasets spanning 10 spatial technologies, eight modalities, and nine tissue types, ranging from thousands of spots to millions of cells, COSIE outperforms existing methods. It resolves tissue structures, enhances noisy measurements, predicts unmeasured modalities, and captures dynamic processes. In human tumors, COSIE identifies invasive subregions linked to clinical outcomes and predicts spatial gene expression in TCGA samples using only histology images. By transforming fragmented data into comprehensive spatial maps, COSIE advances computationally complete spatial omics and the creation of digital tissue twins for biomedicine.

## Introduction

Understanding the molecular and cellular complexity of tissues requires capturing multiple layers of biological information, such as epigenome, transcriptome, proteome, and metabolome, within their native spatial context. Multimodal spatial omics has emerged as a transformative framework to meet this need, offering integrated views of tissue organization and cell states^1^. While early platforms are limited to profiling each modality on separate tissue sections, recent advances have made it possible to simultaneously profile multiple omics layers within the same section, preserving critical cross-modality relationships essential for decoding cellular phenotypes and tissue architecture^2–10^. Despite these technical advances, current experimental approaches remain fundamentally constrained, where only a small number of omics modalities can be simultaneously profiled, and efforts to increase multiplexity often compromise data quality. As a result, multimodal spatial omics data are often fragmented, incomplete, and noisy, which hinders downstream interpretation and biological discovery.

Overcoming these limitations requires not only improved experimental technologies, but also more powerful computational solutions. Yet existing computational methods for multimodal integration remain inadequate: many are narrowly tailored or overfitted to specific modality combinations, lack scalability to cellular or subcellular resolution^11^, or fail to account for spatial context, treating inherently structured tissue data as if they were dissociated single cells^12–18^. Therefore, there is a pressing need for a unified computational framework that can integrate heterogeneous spatial omics data, predict missing modalities, enhance quality of measurements, and scale across tissue types, platforms, and datasets that involve millions of cells.

Here, we introduce COSIE (Within- and **C**r**O**ss-subject **S**patial multimodal **I**ntegration, prediction and **E**nhancement), a virtual machine that computationally synthesizes tissue sections profiled with diverse modalities into a unified latent space, enabling virtual prediction of unmeasured omics layers, enhancement of noisy measurements, and harmonization of incomplete datasets across tissue sections, individuals, and platforms. Importantly, COSIE accommodates a wide range of input types, including epigenome, transcriptome, proteome, metabolome, and histology, and is compatible with both spot-level and single-cell resolution spatial omics platforms. We demonstrate COSIE’s broad applicability by analyzing 12 datasets spanning 10 mainstream spatial profiling technologies, eight different modalities, and nine tissue types, including large-scale datasets encompassing millions of cells (**Supplementary Table 1**). Moreover, COSIE-predicted molecular profiles from hematoxylin and eosin (H&E) stained histology images faithfully reveal fine-grained and functionally relevant spatial patterns, and enable the identification of survival-associated genes in The Cancer Genome Atlas (TCGA) lung cancer samples.

By integrating diverse data types into coherent and biologically meaningful representations, COSIE establishes a computational foundation for the development of digital twins of solid tissues. These high-resolution virtual models capture molecular and spatial complexity across individuals and cohorts. Together, these results position COSIE as a general-purpose, scalable, and biologically informative framework that meets key experimental and analytical challenges in multimodal spatial omics, and supports the vision of computationally complete models of virtual cells and tissues^19^.

## Results

### Overview of COSIE

COSIE is a unified and scalable framework for multimodal spatial omics analysis, capable of integrating diverse imaging and molecular modalities, including H&E-stained histology images, chromatin accessibility, histone modifications, transcriptomics, proteomics, and metabolomics, across tissue sections and subjects with heterogeneous and incomplete modality coverage (**Fig. 1a**). Acting as a virtual machine for spatial omics completion, COSIE aims to computationally generate a full set of omics modalities for each tissue section, even when only a subset of modalities is originally measured. COSIE achieves this by embedding all sections into a shared latent space, enabling joint representation learning, and harmonization across heterogeneous spatial omics datasets. The COSIE framework comprises four components (**Fig. 1b and Methods**): (1) a preparatory stage that extracts hierarchical histological features from sections containing H&E images using the histology foundation model, UNI^20^, and constructs cross-section linkages, including strong linkages from shared modalities and weak linkages from biologically related but distinct ones (e.g., RNA and protein, ATAC and RNA); (2) modality-specific encoding using graph autoencoders to learn low-dimensional and biologically meaningful representations for each modality; (3) intra-section multimodal integration via contrastive learning and predictive learning to reinforce shared signals and predict missing modality embeddings; and (4) cross-section harmonization using structure-aware representation learning and triplet loss. Within the fully integrated embedding space, COSIE enables virtual prediction of missing modalities by aggregating molecular profiles from similar cells across sections. This approach helps overcome data sparsity due to the limited sensitivities of current spatial omics platforms^21,22^. By leveraging a subgraph-based training strategy, COSIE scales efficiently to large datasets containing millions of cells. These capabilities establish COSIE as a versatile computational framework for unified, scalable, and modality-complete spatial omics analysis.

**Fig. 1.**
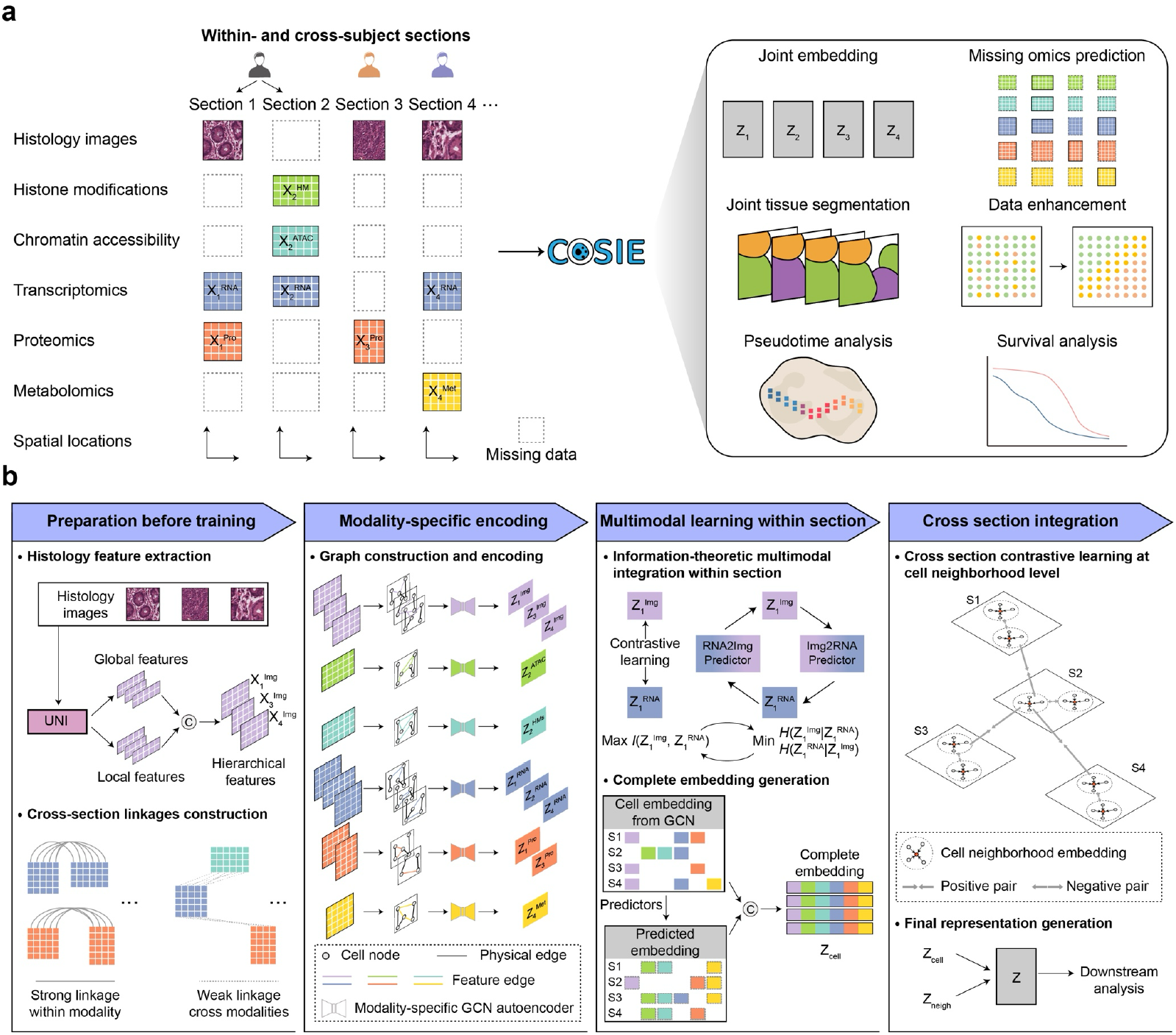
Overview of COSIE for multimodal spatial omics integration, prediction, and enhancement. **a**, COSIE integrates multimodal spatial omics data across multiple tissue sections either within- or cross-subjects, where each section may contain a subset of modalities, including histology images, histone modifications, chromatin accessibility, transcriptomics, proteomics, and metabolomics, along with spatial location. COSIE performs joint embedding and multimodal data completion across all sections, enabling a variety of downstream tasks. **b**, COSIE workflow. (1) If histology images are available, COSIE extracts both global and local features using a hierarchical strategy based on UNI, which are used as inputs for the histology modality. Cross-section linkages are established by leveraging both shared-modality correspondences and biologically relevant cross-modality relationships (RNA–protein, RNA–ATAC, etc.), guiding subsequent integration across sections. (2) For each input data matrix, a graph is constructed by considering both spatial proximity and feature similarity. Modality-specific graph autoencoders are then applied to extract latent representations. (3) Within each section, COSIE performs information–theoretic multimodal integration by jointly optimizing contrastive learning and bi-directional predictors (e.g., RNA2Img and Img2RNA, I(·,·) represents mutual information and H(·|·) denotes conditional entropy), enabling both robust multimodal integration and prediction of missing modality embeddings. (4) COSIE integrates heterogeneous and incomplete sections by linking cells through pre-established linkages at the cell-neighborhood level. Final cell representations are derived by combining both cell-level and neighborhood-level embeddings.

### Revealing complex architecture in human periodontal and atherosclerotic tissues

To evaluate the performance of COSIE, we applied it to a diverse collection of datasets encompassing a wide range of spatial profiling technologies, modalities, and spatial resolutions. We first tested COSIE in a within-subject setting using a human periodontal tissue comprising two immediately adjacent sections (**Fig. 2a**). Section 1 was profiled using H&E and 10x Visium HD, and Section 2 using CODEX and 10x Visium HD. Both Visium HD datasets were processed at an 8 × 8 µm^2^ resolution. To integrate modalities in Section 2, we registered the Visium HD and CODEX data to the same coordinate space and extracted matched protein signals in the corresponding 8 × 8 µm^2^ cell bins in Visium HD to ensure consistent spatial resolution across transcriptomic and proteomic data. Cell phenotypes derived from CODEX data in Section 2 are shown in **Fig. 2b**. We compared COSIE with StabMap^12^, a leading method for single-cell mosaic integration and, to our knowledge, the only available method capable of jointly analyzing more than two modalities, including H&E, while remaining computationally feasible for large-scale datasets. Both COSIE and StabMap jointly integrated the two sections and revealed coherent spatial structures aligned with the cell phenotypes inferred from CODEX (**Fig. 2b,c**). K-means clustering of COSIE embeddings identified biologically meaningful clusters: clusters 0, 3, 4, 6, 9, and 10 corresponded to epithelial cells; cluster 1 to endothelial cells; clusters 2, 5, and 8 to fibroblasts; and cluster 11 to immune cell populations. Although StabMap also captured the broad tissue architecture, COSIE identified more immune cells, better matching the immune cell distribution in the CODEX-based phenotypes (**Fig. 2b,c**, where cluster 11 corresponds to immune cells in both methods).

**Fig. 2.**
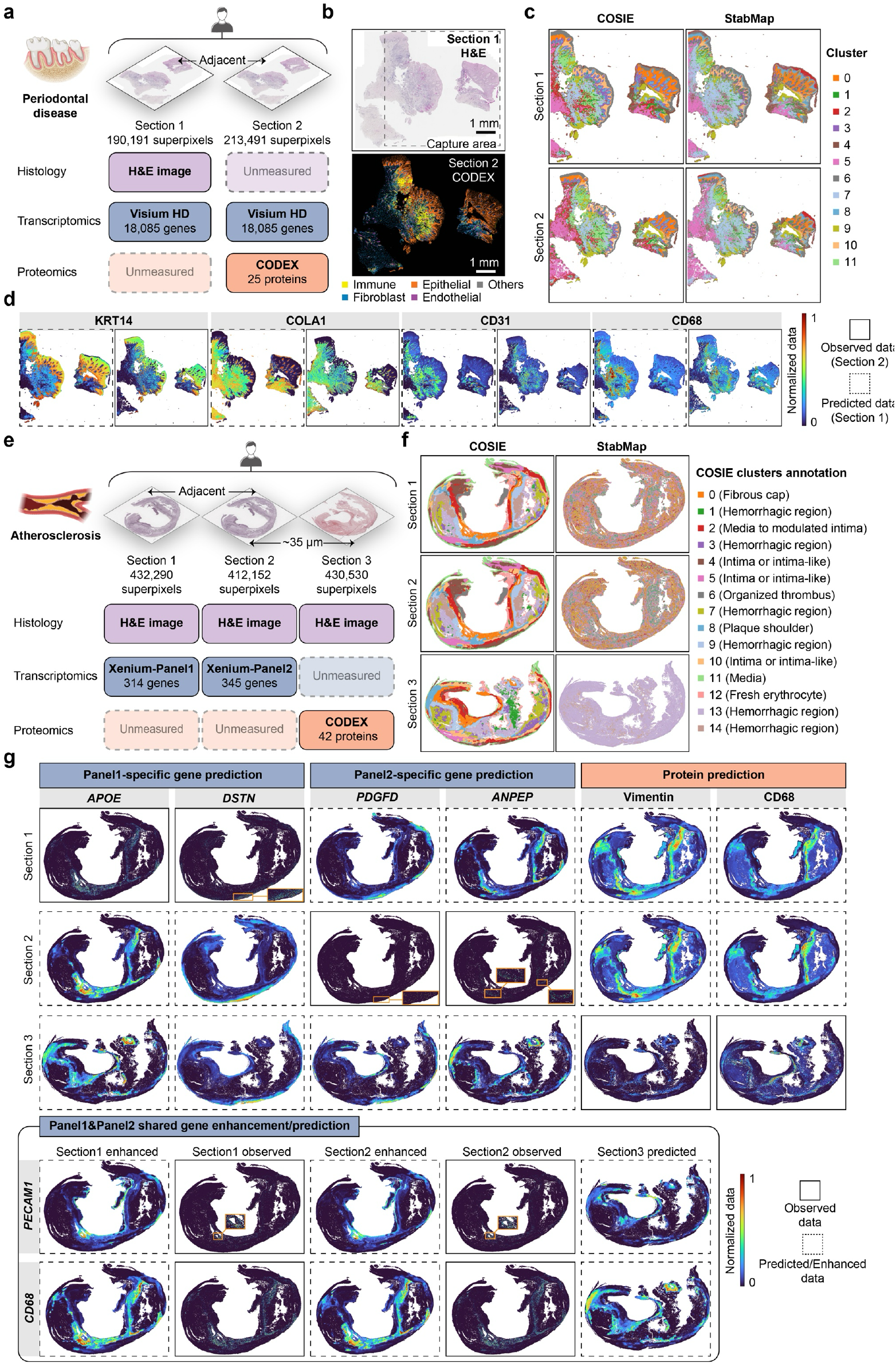
Analysis of spatial transcriptomics–proteomics data in human periodontal and atherosclerotic tissues. **a-d**, Within-subject analysis of a human periodontal disease dataset comprising two adjacent tissue sections. **a**, Experimental design of human periodontal disease dataset. Section 1 includes paired H&E staining and 10x Visium HD spatial transcriptomics. Section 2 includes paired 10x Visium HD transcriptomics and CODEX spatial proteomics. Created using BioRender.com. **b**, H&E image in Section 1 (top) and cell phenotypes based on CODEX data in Section 2 (bottom). **c**, Joint segmentation results from COSIE and StabMap on the periodontal dataset. **d**, Protein prediction for Section 1 using COSIE, with Section 2 serving as the reference. For each protein, the dashed boxes (left) represent predicted protein abundance in Section 1, and the solid boxes (right) show observed CODEX measurements in Section 2. To reduce the impact of extreme outliers in visualization, observed expression values for each protein were truncated at the 99th percentile of non-zero values. All values were normalized to [0, 1]. **e-g**, Within-subject analysis of a human atherosclerosis dataset comprising three tissue sections. **e**, Experimental design of human atherosclerotic plaque dataset. Sections 1 and 2 were immediately adjacent and profiled using 10x Xenium with different gene panels (panel 1: 314 genes; panel 2: 345 genes; 111 genes shared), while Section 3 was profiled using CODEX to measure 42 proteins. All sections include H&E staining. Created using BioRender.com. **f**, Joint segmentation results from COSIE and StabMap on the atherosclerosis dataset. Cluster annotations for the COSIE results are shown on the right. **g**, Predicted and enhanced expression patterns from COSIE. Top panels show prediction of panel-specific genes and proteins across three sections. Bottom panels show prediction and enhancement of genes shared between Xenium panel 1 and panel 2, where enhanced and observed expression in Sections 1 and 2, along with predicted expression in Section 3 (right) are shown. Solid boxes denote observed data, and dashed outlines denote predicted or enhanced values. To reduce the impact of extreme outliers in visualization, observed expression values for each gene were truncated at the 99th percentile of non-zero values. Several key regions were magnified for clearer illustration due to the high sparsity of the observed gene expression data. All values were normalized to [0, 1].

Using COSIE’s learned embedding, we next predicted missing protein expression in Section 1 by leveraging Section 2’s directly measured CODEX data as a reference. COSIE-predicted protein abundance patterns closely matched those from CODEX in Section 2 (**Fig. 2d**). For example, KRT14, a canonical epithelial marker^23^, exhibited a strong and spatially localized expression pattern consistent with epithelial cell distribution. The fibroblast marker COLA1^24^ displayed similarly structured spatial pattern compared to the original measurement. Moreover, CD31 and CD68, markers of endothelial and myeloid cells^25,26^, respectively, also closely recapitulated their known spatial localization. **Supplementary Fig. 1** shows additional results. For comparison, we also applied StabMap to predict protein expression using its learned embeddings (**Supplementary Fig. 2**). Both methods performed well and yielded cleaner spatial localization of protein markers.

Encouraged by these results, we next evaluated COSIE in a more challenging scenario: human atherosclerotic plaque tissue profiled across three sections from a patient undergoing carotid endarterectomy (**Fig. 2e**). Two immediately adjacent sections were profiled using 10x Xenium, each with a different target gene panel (314 genes for Section 1, 345 genes for Section 2, with 111 genes shared), and paired H&E images. The third section, which is ∼35 µm apart, was profiled using CODEX, with a paired H&E image from an adjacent slice. All data were harmonized to 8 × 8 µm^2^ superpixels, and transcript/protein signals were aggregated within corresponding pixels. This integration task is more complex than the periodontal case due to larger inter-section distances, greater morphological variation, and limited modality overlap, with H&E being the only shared modality across all three sections. COSIE successfully integrated the three sections using both strong (H&E–H&E, RNA–RNA) and weak linkages (RNA–protein) to embed corresponding tissue regions while preserving anatomical structures (**Fig. 2f**). It resolved fine-grained spatial domains, including the media (cluster 11), organized thrombus (cluster 6), plaque shoulder (cluster 8), fresh erythrocyte (cluster 12), hemorrhagic regions (clusters 1, 3, 7, 9, 13, and 14), fibrous cap (cluster 0), and media to modulated intima zones (cluster 2), as well as intima-like areas (clusters 4, 5, and 10). By contrast, StabMap failed to resolve several of these finer structural regions, highlighting COSIE’s superior capacity to model complex tissue architecture across spatially and molecularly heterogeneous datasets.

We further evaluated COSIE’s prediction and signal enhancement capabilities across the integrated embedding space (**Fig. 2g, Supplementary Figs. 3,4**). COSIE predicted genes and proteins not measured or missing across sections and enhanced shared genes between the two Xenium panels. For example, *APOE*, a marker of macrophages^27^, showed strong spatial patterns in relevant anatomical regions such as the atherosclerotic plaque shoulder and the hemorrhagic zones (cluster 9). Smooth muscle markers *DSTN* and *PDGFD*^28,29^, showed clear expected localization in the media (cluster 11) and intima-like areas. For the shared genes between two Xenium panels like *PECAM1*^25^, COSIE enhanced expression patterns were more spatially resolved and matched between the two panels. In particular, *PECAM1* was enriched in fibrous cap (cluster 0), with consistent signals across all three sections. Moreover, for CD68, COSIE accurately predicted both protein- and gene-level expression, with spatial patterns concordant across sections and modalities. These results underscore COSIE’s ability to build a unified latent space that supports robust integration, prediction, enhancement, and precise tissue mapping from heterogeneous spatial omics data to enable virtual tissue construction.

**Fig. 3.**
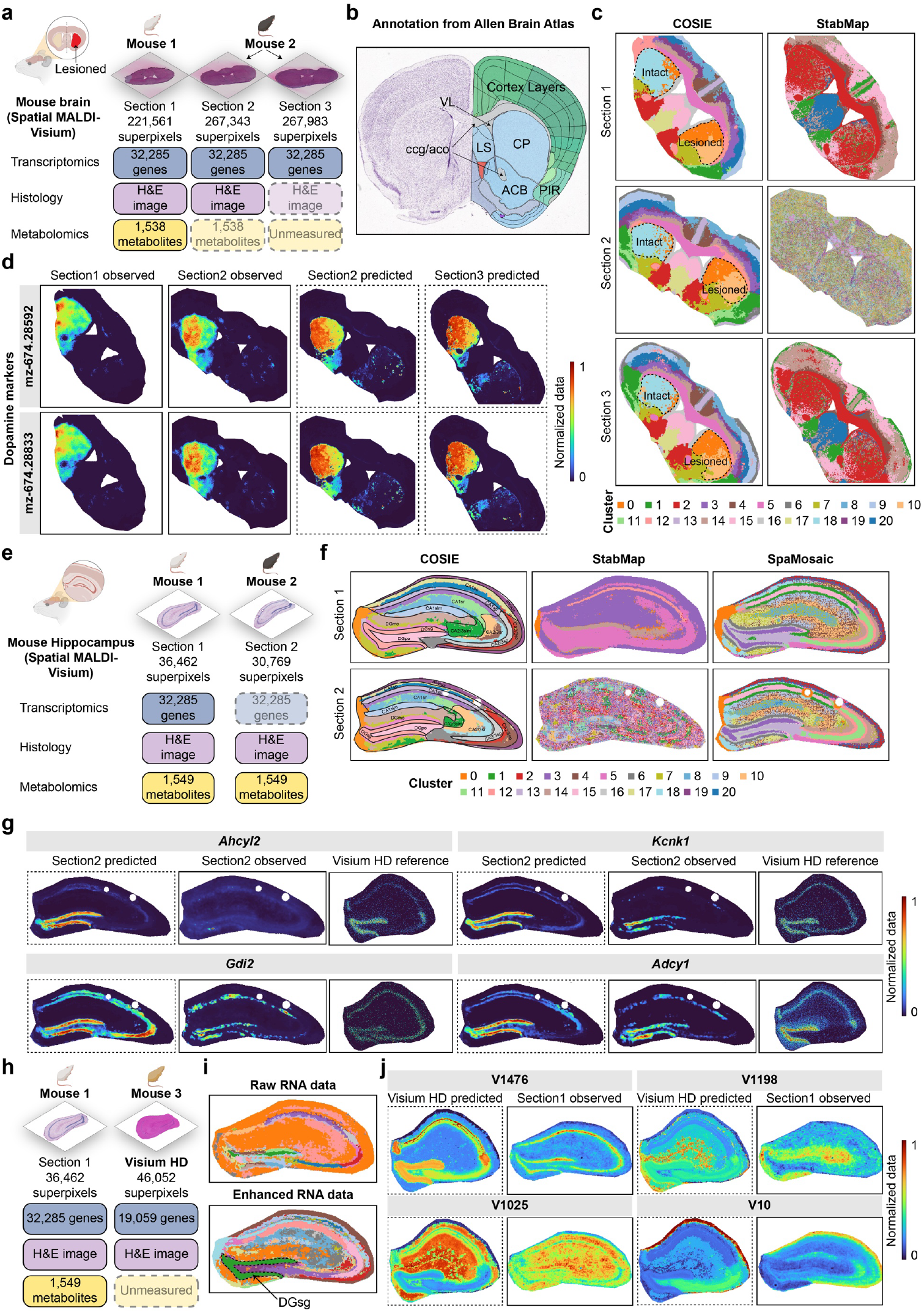
Analysis of two spatial transcriptomics–metabolomics datasets in mouse brains. **a-d**, Application to a spatial MALDI-Visium dataset from adult mouse brains with 6-OHDA-induced lesion. **a**, Experimental design showing integration across three brain sections from two mice. Section 1 contains paired RNA, H&E, and metabolomics data; Section 2 includes RNA and H&E, with metabolomics masked; Section 3 includes RNA only, with H&E masked. Created using BioRender.com. **b**, Anatomical reference from the Allen Brain Atlas. **c**, Joint segmentation results from COSIE and StabMap on the mouse brain dataset. **d**, COSIE-based prediction of metabolite abundance. For each metabolite, shown from left to right are the observed data in Sections 1 and 2, and predicted values in Sections 2 and 3. All values were normalized to [0, 1]. **e-g**, Application to the second spatial MALDI-Visium dataset from mouse hippocampus. **e**, Experimental design showing integration across two hippocampal sections from different mice. RNA data were masked in Section 2. Created using BioRender.com. **f**, Joint segmentation results from COSIE, StabMap and SpaMosaic on the hippocampus dataset. **g**, RNA prediction results for Section 2. For each gene, shown from left to right are COSIE-predicted expression, observed data in Section 2, and the corresponding reference from a high-quality Visium HD dataset. To reduce the impact of extreme outliers in visualization, observed expression values for each gene in the VisiumHD dataset were truncated at the 99th percentile of non-zero values. All data were normalized to [0, 1]. **h-j**, Integration of the spatial MALDI-Visium hippocampal Section 1 with Visium HD simultaneously enhances RNA quality in Section 1 and predicts metabolomics data for the Visium HD section. **h**, Experimental design showing the integration of MALDI Section 1 with Visium HD section. Created using BioRender.com. **i**, RNA-based segmentation results of Section 1(21 clusters). Top: segmentation using original RNA data; bottom: segmentation using enhanced RNA after integration with Visium HD. **j**, COSIE-based metabolites prediction for the Visium HD section. For each metabolite, dashed boxes indicate predicted data of Visium HD, and solid boxes show observed data in MALDI Section 1. All data were normalized to [0, 1].

**Fig. 4.**
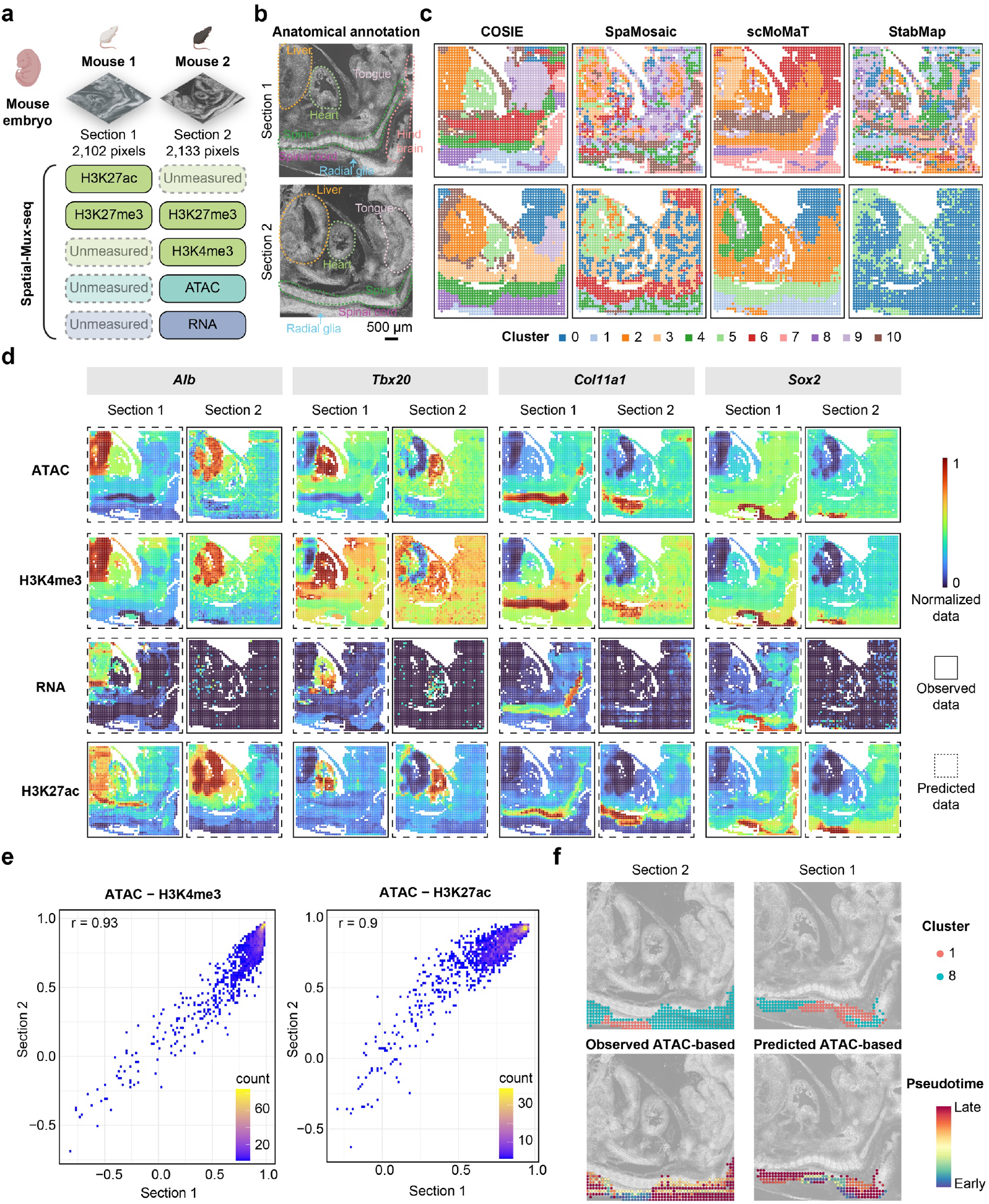
Analysis of spatial epigenomics–transcriptomics across five modalities in mouse embryo. **a**, Experimental design showing integration across two sections from two mouse embryos. Section 1 contains two modalities: H3K27me3 and H3K27ac; Section 2 contains four modalities: ATAC, H3K27me3, H3K4me3, and RNA. Created using BioRender.com. **b**, Anatomical annotation for two sections. The radial glia region is annotated based on marker expression. **c**, Joint segmentation results from COSIE, SpaMosaic, scMoMaT, and StabMap. **d**, Visualization of COSIE-predicted and observed data for representative genes (*Alb, Tbx20, Col11a1*, and *Sox2*) across the two sections. Each row corresponds to a modality (ATAC, H3K4me3, RNA, and H3K27ac), and each column corresponds to a gene. Dashed boxes indicate predicted data, and solid boxes show observed values. **e**, Cross-section consistency of gene-wise cross-modality correlations. Each panel shows the correlation of gene-wise Pearson correlation coefficients between ATAC and a histone modification (H3K4me3, left; H3K27ac, right), computed separately within Section 1 and Section 2. COSIE-predicted ATAC and H3K4me3 data were used in Section 1, and COSIE-predicted H3K27ac data were used in Section 2. High concordance indicates that COSIE preserves cross-modality regulatory relationships across sections. Color indicates gene density. **f**, Pseudotime inference based on observed ATAC (Section 2) and predicted ATAC (Section 1) data reveals consistent spatial developmental trajectories. Two representative clusters (1 and 8), corresponding to radial glia and differentiated neurons in the spinal cord region, are highlighted (top panel). The bottom panel shows pixels colored by inferred pseudo-time values.

### Uncovering fine-grained structures in mouse brain

To assess COSIE’s performance on additional omics types, we next applied it to spatial MALDI-Visium datasets combining 10x Visium and MALDI mass spectrometry imaging (MALDI– MSI), which jointly profile H&E staining, metabolomics, and transcriptomics on the same tissue section. We first analyzed an adult mouse brain dataset^8^ (8 weeks old) from a Parkinson’s disease model induced by 6-hydroxydopamine (6-OHDA), a neurotoxin that selectively depletes dopaminergic neurons (**Fig. 3a**). As reported in the original study^8^, dopamine and related metabolites were strongly enriched in the intact striatum, clearly delineating hemispheric asymmetry. By contrast, transcriptomic and H&E data alone lacked sufficient contrast to resolve this spatial distinction. Three sections were analyzed: Section 1 from one mouse, and Sections 2 and 3 from another. Sections 1 and 2 contained paired H&E, metabolomics and transcriptomics data, while Section 3 included only H&E and transcriptomics. To ensure the same spatial resolution, Visium and MALDI data were first aligned using SpaMTP^30^, and then enhanced to 8 × 8 µm^2^ resolution with iStar^31^. To simulate real-world multimodal incompleteness, we masked metabolomics data in Section 2 and H&E in Section 3. The Allen Brain Atlas annotation for P56 mouse brain coronal section was used as the anatomical reference (**Fig. 3b**). COSIE successfully leveraged the available metabolomics data in Section 1 to resolve lesion-intact hemispheric asymmetry across all sections (**Fig. 3c**). Notably, COSIE distinguished the lesioned (clusters 0 and 10) and intact caudoputamen (CP) (cluster 18), and accurately delineated laminar structures (clusters 3, 6, 8, 9, 14, 19, and 20), and other brain regions, such as lateral septal complex (LS, cluster 15), lateral ventricle (VL, cluster 16), corpus callosum genu/the anterior cortical nucleus of the amygdala (ccg/aco, clusters 5 and 12), piriform area of the cortex (PIR, cluster 1), nucleus accumbens (ACB, clusters 2 and 7), and cingulate cortex (cluster 4). COSIE also predicted dopamine localization in Sections 2 and 3 where metabolites were masked, recapitulating its enrichment in the intact hemisphere (**Fig. 3d**), a key hallmark of 6-OHDA-induced lesions, whereas StabMap failed to resolve this pattern (**Fig. 3c, Supplementary Fig. 5**).

We next assessed COSIE’s ability to resolve fine-grained brain structures in a second spatial MALDI-Visium dataset from mouse hippocampus. Two coronal sections from different mice were profiled using the MISO^32^ described protocol, with paired H&E, metabolomics, and transcriptomics data (**Fig. 3e**). As before, all modalities were enhanced to 8 × 8 µm^2^ resolution. To mimic an incomplete data setting, RNA was masked in Section 2. We compared COSIE with both StabMap and SpaMosaic^11^, a method applicable to moderately sized datasets. COSIE produced anatomically coherent segmentations that aligned closely with known hippocampal architecture (**Fig. 3f**), including the dentate gyrus stratum granulosum (DGsg, cluster 12), polymorph layer (DGpo, cluster 15), and stratum lacunosum-moleculare of CA1 (CA1slm, cluster 9). In contrast, competing methods yielded noisier and less consistent segmentations. For example, COSIE clearly separated the stratum oriens (CA1so) and stratum radiatum (CA1sr) into distinct clusters, whereas SpaMosaic merged them into shared regions. Moreover, COSIE delineated the CA2/3slm region as a distinct structure, while SpaMosaic grouped this area with clusters from CA1sr and CA2/3sr, failing to resolve finer anatomical boundaries. In addition, COSIE’s predicted gene expression in Section 2 closely matched spatial patterns in an independent high-quality Visium HD dataset obtained from a different mouse profiled by 10x Genomics (**Fig. 3g, Supplementary Fig. 6**), demonstrating accurate signal recovery despite the original transcriptomics data’s suboptimal quality.

Finally, we integrated the MALDI-Visium data in Section 1 with the Visium HD dataset to simultaneously enhance gene expression in Section 1 and predict metabolite profiles in the Visium HD section (**Fig. 3h**). COSIE substantially improved gene expression patterns, which enabled clearer segmentation of hippocampal structures such as the DGsg (**Fig. 3i, Supplementary Fig. 7**). Metabolite prediction in the Visium HD section also closely mirrored the original MALDI measurements in Section 1 (**Fig. 3j**). As multimodal spatial omics data grow in complexity, uneven data quality across modalities becomes a major challenge. COSIE offers a unified computational solution to predict missing modalities and enhance noisy measurements through cross-modal, cross-section learning.

### Recovering developmental trajectory in mouse embryo across five modalities

To evaluate COSIE’s ability to integrate highly heterogeneous modalities, we applied it to the Spatial-Mux-seq dataset^10^, which spans five distinct modalities across two E13 mouse embryos, profiled at 50 µm spatial resolution (**Fig. 4a**). Section 1 from one embryo includes H3K27ac and H3K27me3 histone modifications, while Section 2, obtained from another embryo, contains ATAC, RNA, H3K27me3, and H3K4me3. This setting presents substantial challenges due to modality heterogeneity and inter-subject variability. COSIE leveraged the shared H3K27me3 modality (strong linkage) and additional cross-modality connections (weak linkages) to construct a unified latent space. As shown in **Fig. 4b,c**, COSIE accurately segmented major tissue structures and assigned anatomically corresponding regions to the same clusters across sections; for example, the liver (cluster 2), heart (cluster 5), tongue (cluster 9), spine (cluster 4), spinal cord (cluster 8), and radial glia (cluster 1). Notably, the hindbrain, present only in Section 1, was correctly identified as a distinct region (cluster 7). As all modalities in this dataset are omics-based, scMoMaT^13^ was included for comparison. As shown in **Fig. 4b,c**, competing methods produced noisier or less coherent segmentation results.

We next predicted missing modalities, including ATAC, H3K4me3, and RNA for Section 1, and H3K27ac for Section 2. We found COSIE’s outputs aligned well with biological expectations (**Fig. 4d**). For example, *Alb* (a canonical liver marker^33,34^) and *Sox2* (enriched in radial glia^35^) showed accurate and enhanced spatial patterns in the predicted data, even when the original RNA signals in Section 2 were weak. Additional examples are provided in **Supplementary Fig. 8**. To assess whether COSIE predictions preserve inter-modality relationships, we computed gene-wise correlations based on the completed data matrices. As a representative example, for each gene, we calculated the Pearson correlation between its ATAC and H3K4me3 profiles across pixels in Section 1 (predicted data-based) and Section 2 (observed data-based), respectively. We then computed the correlation between these two sets of gene-wise correlation values. As shown in **Fig. 4e**, the result shows strong consistency (r=0.93), indicating that COSIE faithfully predicts not only individual modalities, but also the underlying inter-modality relationships that generalizes across subjects. Similar consistency in correlation patterns was observed across other modality pairs (**Supplementary Fig. 9**), highlighting COSIE’s ability to predict biologically meaningful multimodal interactions across subjects.

Finally, we performed pseudotime analysis to assess COSIE’s ability to preserve dynamic spatiotemporal relationship during development. We focused on spinal cord, a region known to exhibit a radial glia-to-neuron trajectory^10,36^, and selected ATAC as the input modality given its established role in capturing regulatory potential and cell state transitions. In Section 2, where ATAC was directly profiled, COSIE-identified clusters 1 and 8 corresponded to radial glia and spinal cord, respectively. Pseudotime inference revealed a smooth spatial gradient, with central cells showing lower pseudotime values, indicative of radial glia, while peripheral cells exhibited more advanced pseudotime, consistent with the known trajectory from radial glia to postmitotic neurons^10,36^ (**Fig. 4f**, left panel). Notably, this trajectory closely aligned with COSIE’s segmentation. To assess whether we can obtain similar trajectories from predicted data, we repeated the analysis in Section 1, where ATAC was entirely missing. Remarkably, pseudotime inference based on COSIE-predicted ATAC data revealed a comparable spatial continuum (**Fig. 4f**, right panel), consistent with the expected differentiation axis. These findings demonstrate that COSIE predictions preserve biologically meaningful dynamics and enable accurate downstream inference.

### Resolving tumor heterogeneity in human gastric cancer

Having established COSIE’s effectiveness on non-malignant tissues, we next applied it to gastric tumor samples, which exhibit greater molecular and structural heterogeneity compared with the previously analyzed data. As shown in **Fig. 5a**, two sections obtained from different patients were profiled. Section 1 included Visium spatial transcriptomics and H&E staining; Section 2 included COMET spatial proteomics (19 proteins) and H&E staining from an adjacent section. The only shared modality in these two sections is H&E, which is common in real-world studies. For Section 1, we applied iStar to enhance its gene expression resolution to 8 × 8 µm^2^. For Section 2, its protein abundance was quantified at the 8 × 8 µm^2^ resolution by aligning COMET to the H&E-derived superpixels. Spot-level pathologist annotations for both sections are shown in **Fig. 5b** (by I.L. and L.M.S.S.). COSIE produced anatomically coherent joint segmentations across the two tumor sections and accurately delineated most key tissue regions (**Fig. 5c**). Clusters 4 and 14 corresponded to gastric mucosa, which is rich in glandular and immune cells critical for maintaining mucosal barrier function and host defense. Clusters 6 and 10 marked gastric surface epithelium and mucin-producing regions. The smooth muscle region was captured by clusters 0, 8, 11, 12, 15, 16, and 17, reflecting the organized architecture of the muscularis layer essential for gastric motility. Importantly, tumor-associated regions were identified as clusters 2, 5, and 7, which displayed distinct and morphologically irregular patterns that are characteristic of malignant compartments. These results highlight COSIE’s ability to resolve heterogeneous tumor microenvironments across different tumors and modalities.

**Fig. 5.**
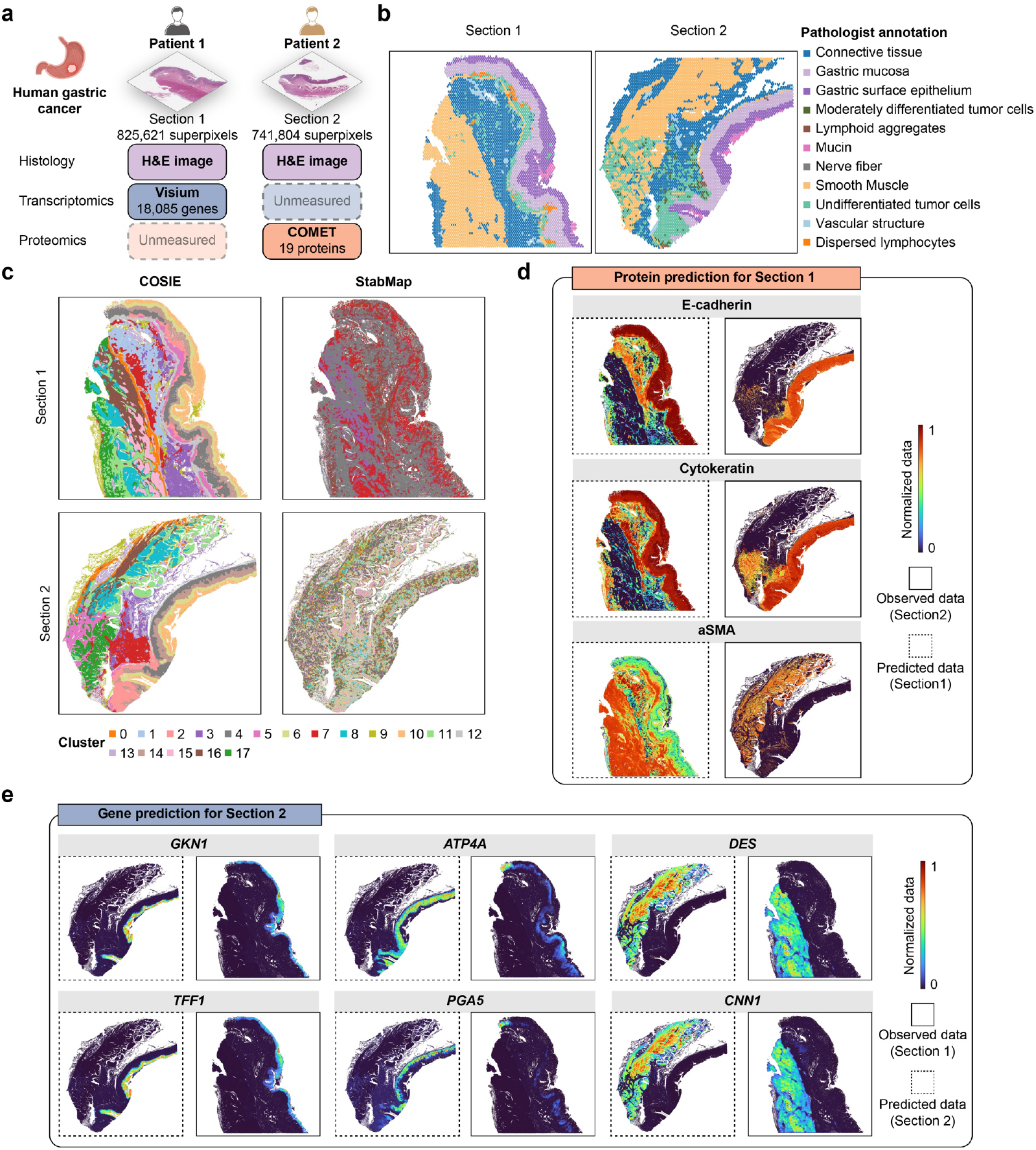
Analysis of human gastric cancer data. **a**, Experimental design of the human gastric cancer dataset. Two sections were generated from different patients. Section 1 included paired H&E and Visium data. Section 2 contained COMET protein and H&E from an adjacent section. Created using BioRender.com. **b**, Pathologist annotations of two gastric cancer sections. The annotation for Section 2 was derived from an adjacent Visium section. **c**, Joint segmentation results from COSIE and StabMap. **d**, COSIE-based protein prediction for Section 1. Dashed boxes indicate predicted data, and solid boxes show observed COMET data. All data were normalized to [0, 1]. **e**, COSIE-based RNA prediction for Section 2. Dashed boxes show predicted expression in Section 2, and solid boxes show observed RNA measurements. All data were normalized to [0, 1].

We next predicted protein abundance in Section 1 and gene expression in Section 2. This represents a particularly challenging scenario, as the two sections are from different cancer patients and the only shared information is the H&E image. Despite the absence of matched omics modalities, COSIE effectively leveraged cross-subject and cross-modality relationships to predict spatially meaningful patterns (**Fig. 5d**). For example, for Section 1, the predicted abundance for E-cadherin was enriched in gastric mucosa and surface epithelium, consistent with its role in epithelial cell adhesion^37^. Cytokeratin was broadly predicted across the epithelial cells located in the mucosa and undifferentiated tumor regions, reflecting its retained expression in poorly differentiated epithelial-derived cancer cells^38^. α-Smooth muscle actin (αSMA) showed enrichment in smooth muscle regions, further validating the biological accuracy of COSIE’s predictions. For gene expression prediction in Section 2, COSIE predictions are consistent with known tissue localization (**Fig. 5e**). *GKN1* and *TFF1*, markers of surface epithelium involved in mucosal protection and differentiation^39,40^, were highly expressed along the luminal surface, closely matching the observed data. *ATP4A* and *PGA5*^41,42^, involved in acid secretion and digestive enzyme production, showed strong signals in mucosa region. Smooth muscle markers *DES* and *CNN1*^43,44^, were selectively expressed in the muscularis. Additional prediction results are shown in **Supplementary Fig. 10**. Together, these results demonstrate COSIE’s robustness in predicting multimodal spatial landscapes across highly heterogeneous tumor tissues.

### Constructing virtual tissue model in human lung cancer

To further evaluate COSIE’s scalability and robustness in disease contexts, we applied it to lung adenocarcinoma (LUAD), a cancer type characterized by marked molecular and structural heterogeneity. As shown in **Fig. 6a**, four LUAD sections from four different patients were profiled using 10x Visium spatial gene and protein platform. To simulate incomplete data scenarios, we masked certain selected modalities: protein data in Sections 1 and 2, gene expression in Section 3, and all but H&E staining in Section 4. Both gene and protein expression data were enhanced to 8 × 8 µm^2^ resolution with iStar. These datasets span over three million superpixels, posing both scale and heterogeneity challenge that, to our knowledge, no existing methods besides COSIE can address. COSIE generated coherent segmentations across the four sections, capturing both normal and cancer-associated tissue structures with strong concordance to expert annotations (**Fig. 6b,c**, annotations by L.G.). It successfully revealed fine-grained tissue features, such as bronchus (cluster 10), lymphoid aggregates (cluster 18), fibrous region (cluster 20), and vascular structure (cluster 21). Notably, macrophage-enriched regions (cluster 19) were detected in Section 4 despite being absent from the original spot-level pathologist annotations (**Fig. 6b**). A second round of pathologist review confirmed the presence of macrophages in COSIE-identified regions (by L.G.), suggesting COSIE’s capability to support fine-grained annotation and reveal subtle tissue features missed by manual review.

**Fig. 6.**
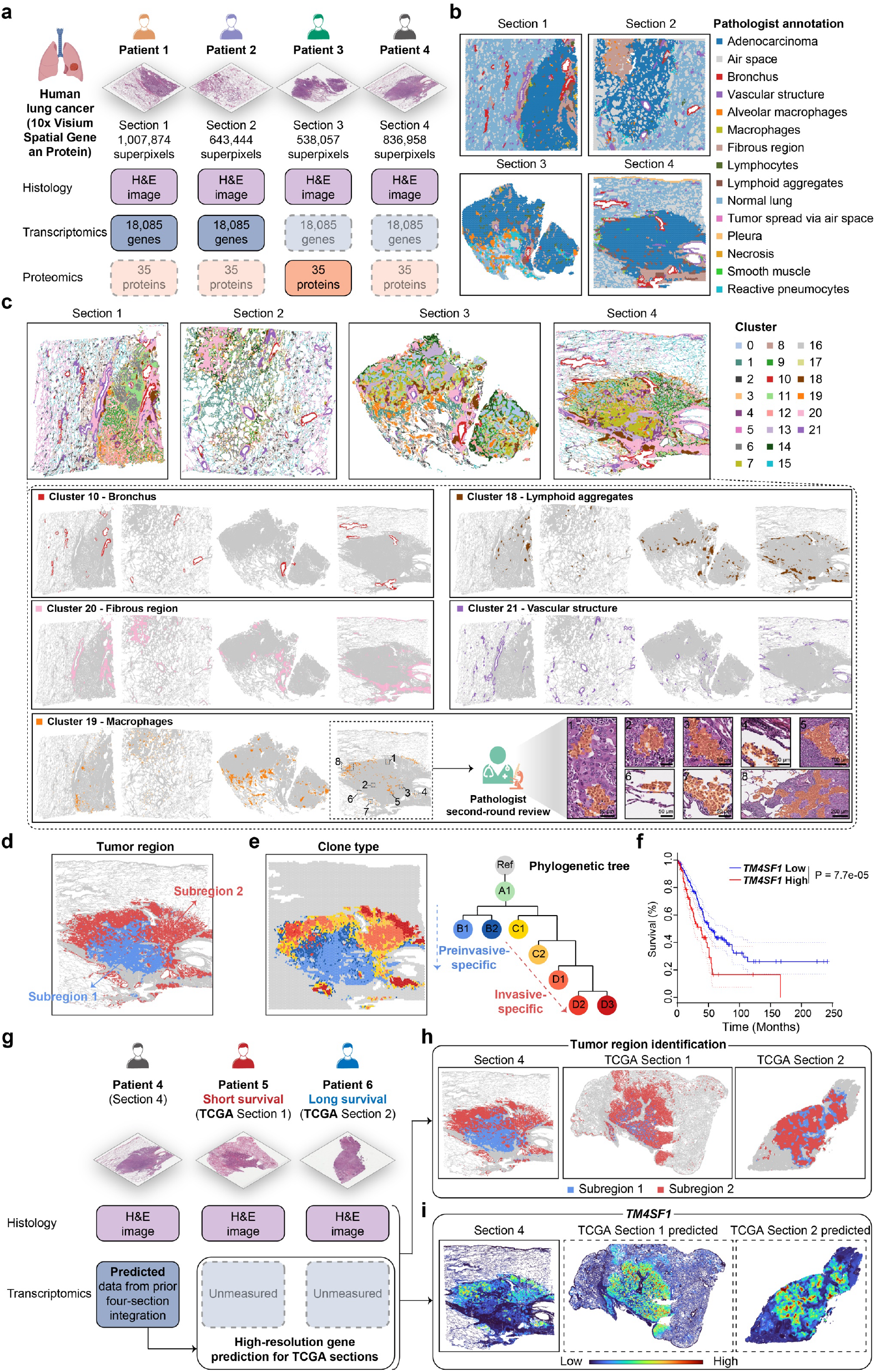
Analysis of human lung cancer data. **a-f**, Integration of human lung cancer data across four sections. **a**, Experimental design of the human lung cancer dataset. Four sections were obtained from four different patients, and profiled using 10x Visium spatial gene and protein platform, generating paired H&E, RNA, and protein data. To simulate an incomplete data scenario, protein data were masked in Sections 1 and 2, RNA data were masked in Section 3, and only the H&E image was retained in Section 4. In total, the dataset contains over 3 million single-cell resolution superpixels. Created using BioRender.com. **b**, Pathologist annotations of four lung cancer sections. **c**, Joint segmentation results from COSIE. For visualization, representative structures were highlighted with their assigned cluster colors, and all other regions are shown in gray. A second-round pathologist review was conducted for cluster 19 to confirm the COSIE-identified macrophages-rich superpixels. **d**, Two tumor subregions identified by COSIE of Section 4. **e**, Clone type identification of Section 4. Left: spatial distribution of clone types identified based on observed spot-level gene expression data, where each spot is colored by its assigned clone identity inferred from CNV profiles. Right: reconstructed phylogenetic tree structure, highlighting the clonal evolution from early lesions to preinvasive (B1, B2) and invasive clones (C1-C2, D1-D3). **f**, Kaplan–Meier curves showing overall survival of LUAD patients stratified by *TM4SF1*. Survival analysis was performed on the TCGA-LUAD cohort (n = 479) using GEPIA2, with a 75% cutoff to define high and low groups. Statistical significance was assessed using the log-rank test. Dotted lines indicate the 95% confidence interval. **g–i**, Integration of Section 4’s H&E image and predicted RNA data with two TCGA H&E-only sections using COSIE. **g**, Experimental design illustrating COSIE-based integration of Section 4 with two TCGA sections. Section 4 used H&E image and predicted RNA data from prior four-section integration. The two TCGA samples, selected based on *TM4SF1* bulk RNA-seq levels and survival, include one from a short-term survivor with high *TM4SF1* expression and one from a long-term survivor with low expression. Both samples contain only H&E images. **h**, Tumor region identification in two TCGA sections through integration with Section 4 by COSIE. **i**, Visualization of COSIE-predicted spatial expression of *TM4SF1*. For Section 4, spatial gene expression was predicted from prior four-section integration. For the two TCGA sections, spatial gene expression was predicted using Section 4-predicted gene expression.

Next, we assessed COSIE’s ability for missing modality prediction, focusing on the most challenging scenario in Section 4, where only H&E image is available (**Supplementary Fig. 11**). COSIE accurately predicted gene expression, including *FOXJ1*, a canonical marker of bronchial epithelium^45^, whose predicted localization matched Xenium data derived from an adjacent tissue section. Predicted protein signals also reflected known biology. Quantitative evaluation showed that COSIE preserved RNA–protein concordance, with Pearson correlation between paired modalities remaining high for both observed and COSIE-predicted data (**Supplementary Fig. 12**). Segmentation based on predicted molecular profiles also delineated major tissue structures (**Supplementary Fig. 13**). These findings underscore COSIE’s capability as a virtual machine for computational generation of multimodal spatial omics data.

To explore intratumoral heterogeneity, we analyzed COSIE derived segmentations in Section 4 and identified two distinct tumor subregions by merging spatially coherent clusters based on their proximity in the integrated embedding space using Euclidean distance (**Fig. 6d, Supplementary Fig. 14**). To investigate the mechanisms underlying LUAD aggressiveness, we performed differential gene expression analysis between the two tumor subregions using COSIE-predicted data and identified 200 highly expressed genes for each subregion. Gene set enrichment analysis suggested Subregion 2 is more invasive (**Supplementary Fig. 15**). To validate this finding, we conducted copy number variation (CNV) and pseudotime analyses using the observed gene expression data from Section 4. CNV analysis using SpatialInferCNV^46^ revealed a clear phylogenetic separation between preinvasive and invasive clones, with Subregion 1 enriched for preinvasive cells and Subregion 2 enriched for invasive ones (**Fig. 6e**). Pseudotime analysis further revealed a developmental trajectory consistent with the tumor progression inferred from the phylogenetic tree (**Supplementary Fig. 16**), confirming that Subregion 2 is more invasive.

Next, we investigated whether candidate therapeutic targets could be identified from the genes enriched in Subregion 2. We applied a multi-step filtering strategy emphasizing translational potential. First, we selected genes encoding cell surface proteins based on predicted subcellular location from the Human Protein Atlas^47^, given their accessibility for therapeutic targeting and biomarker development. Among the 26 surface protein-encoding genes, we further performed survival analysis in the TCGA LUAD bulk RNA-seq samples (n = 479) to assess association between gene expression and overall survival for each gene. After correction for multiple testing, *TM4SF1* emerged as the only gene showing a statistically significant association with survival (**Fig. 6f, Supplementary Table 2**), underscoring its clinical relevance. *TM4SF1* is a tetraspanin-like surface protein implicated in cell migration, proliferation, and tumor progression, and has been reported as a marker of cancer stem-like cells and poor prognosis in multiple epithelial malignancies^48,49^. Identifying *TM4SF1* not only highlights a surface-accessible marker linked to tumor aggressiveness, but also provides a promising entry point for therapeutic development in LUAD. These findings demonstrate the utility of COSIE-predicted gene expression in uncovering biologically informative and potentially druggable targets.

Building on these findings, we applied COSIE to integrate Section 4 with LUAD samples from TCGA using only their H&E images (**Fig. 6g**). Section 4, with COSIE-predicted gene expression from the prior integration (**Fig. 6a**), served as the reference. Two TCGA samples were selected based on their *TM4SF1* bulk RNA-seq levels and survival outcomes: one with high expression and short survival, the other with low expression and long survival. COSIE successfully transferred the spatial architecture from Section 4 onto the TCGA samples, identifying key tissue regions such as bronchus, vascular structure, fibrous region, and lymphoid aggregates (**Supplementary Fig. 17**), and resolving two tumor subregions (**Fig. 6h, Supplementary Fig. 18**). These closely matched subregions identified in Section 4 from the prior integration (**Fig. 6d**), highlighting COSIE’s reproducibility. In the short-survival sample, the more invasive subregion (Subregion 2) was enriched at the tumor periphery, while this pattern was weaker in the long-term survival sample. COSIE also predicted high resolution spatial gene expression for the two TCGA samples (**Fig. 6i, Supplementary Fig. 19**). *TM4SF1* again exhibited peripheral enrichment in the short-survival case. Notably, this integration was based solely on H&E, and the reference gene expression in Section 4 was itself predicted, demonstrating COSIE’s ability to infer biologically meaningful structures from incomplete inputs. These capabilities represent a foundational step toward virtual spatial omics.

### Generalizability across tissues and technologies

We further evaluated COSIE’s generalizability across different tissue types and spatial omics technologies. As shown in **Supplementary Fig. 20**, COSIE was applied to human tonsil sections profiled by 10x Visium spatial gene and protein platform. It effectively integrated two sections, consistently identifying all germinal centers (GCs). Predicted molecular profiles data exhibited strong spatial concordance with observed data; for example, PCNA, a canonical GC marker^50^, showed consistent spatial localization between the predicted and measured signals. Gene– protein correlations were also well preserved, further supporting the biological fidelity of COSIE’s predictions. In the diffuse large B-cell lymphoma (DLBCL) dataset (**Supplementary Fig. 21**), COSIE successfully propagated CosMx gene expression from limited field-of-views (FOVs) to the entire tissue core and enabled prediction of CODEX protein expression from H&E alone. Additional analyses shown in **Supplementary Fig. 22** demonstrate COSIE’s effectiveness on mouse spleen data profiled by SPOTS^3^. SpaMosaic also performed well on this dataset. A summary of all datasets analyzed in this study is provided in **Supplementary Table 1**.

## Discussion

In this work, we introduced COSIE, a general-purpose virtual machine designed to advance toward computationally complete spatial omics by integrating, predicting, and enhancing multimodal spatial omics data across tissue sections, individuals, and platforms. COSIE enables unified analysis across datasets with heterogeneous and incomplete modality coverage, addressing one of the key bottlenecks in current spatial biology: the lack of comprehensive, high-quality multimodal omics measurements in individual tissue sections. Unlike existing computational tools that are often restricted to two modalities or require paired observations, COSIE flexibly scales to multiple omics layers, different modality combinations, and variable spatial resolutions. It is also highly efficient; for example, COSIE completed training process for integrating four human LUAD sections encompassing over 3 million single-cell resolution superpixels in 23 minutes on a single NVIDIA A100 GPU with 80 GB memory. By contrast, existing methods such as SpaMosaic are not scalable to large spatial datasets, and StabMap requires substantially longer training times on datasets where it is applicable (**Supplementary Fig. 23**).

We validated COSIE extensively across a diverse panel of real world datasets, spanning eight modalities, including histology, chromatin accessibility, histone modifications (H3K27me3, H3K27ac, H3K4me3), transcriptomics, proteomics, and metabolomics, generated from 10 widely adopted spatial profiling technologies, comprising Visium, Visium HD, Xenium, CosMx, CODEX, COMET, 10x Visium spatial gene and protein platform, SPOTS, Spatial-Mux-seq, and spatial MALDI-Visium. These datasets covered a wide spectrum of tissue types and biological contexts, from mouse brain, embryo, and spleen, to human tonsil, lymph node, periodontal disease, atherosclerotic plaque, gastric cancer, and lung cancer, reflecting a continuum of biological states, from normal physiology to inflammation and malignancy. COSIE seamlessly integrated data from similar and distinct modalities, even when generated by different panels or platforms, and performed consistently across spatial scales, from thousands of low-resolution spots to millions of single-cell resolution superpixels within and across individuals, all processed under a single set of default hyperparameters, without dataset-specific tuning. This robustness demonstrates COSIE’s wide applicability, user-friendliness, and accessibility for the broader research community.

Importantly, COSIE enables not only cross-modality integration but also virtual prediction of unmeasured modalities from limited or single-modality input. Across normal and diseased tissues, COSIE predicted spatially coherent and biologically meaningful molecular profiles. For instance, it predicted intact hemisphere-specific dopamine metabolite in mouse brain, inferred developmental trajectories in mouse embryo through virtual ATAC analysis, and transferred high-resolution gene expression maps to TGCA LUAD samples with only H&E input, using a reference section whose gene expression was itself virtually predicted. This layered, cross-sample, cross-subject propagation of virtual molecular information, building predictions on top of predictions, exemplifies COSIE’s unique capacity to disseminate molecular insights across tissues, individuals, and cohorts. By bridging histology with omics data in a fully computational manner, COSIE offers a powerful framework for augmenting traditional pathology and extending molecular analysis to large-scale archival datasets lacking spatial omics measurements, with strong potential for translational and clinical applications.

Despite these strengths, several limitations remain. COSIE’s performance may degrade in extreme settings where all input modalities are simultaneously sparse or noisy. Additionally, while COSIE provides a unified framework for multimodal spatial omics integration, it currently operates on static snapshots and does not explicitly model temporal dynamics, such as development or disease progression. Future extensions to jointly capture spatiotemporal trajectories and cellular lineage relationships would further expand COSIE’s capabilities.

In summary, COSIE provides a scalable, flexible, and generalizable solution to key limitations in spatial omics. Without COSIE, generating multimodal spatial omics datasets with complete molecular modality coverage at the scale investigated in this paper would be experimentally infeasible. By addressing this otherwise intractable challenge, COSIE reduces experimental cost and burden, enhances data quality, and enables deeper biological discovery. COSIE marks a major step toward computationally complete, high-resolution digital twins of tissues, advancing the broader vision of virtual tissue models for biomedical research and precision medicine.

## Methods

### COSIE framework

### Omics data preprocessing

COSIE is able to integrate diverse omics modalities across multiple tissue sections, including epigenome (ATAC-seq, histone modifications), transcriptome, proteome, and metabolome data. Each modality undergoes a tailored preprocessing pipeline reflecting its unique characteristics. For ATAC and histone modification data, gene activity scores are first computed as proxies for regulatory activity, followed by library-size normalization and log transformation. For transcriptomic data, the top 3,000 highly variable genes (HVGs) are selected, then raw counts are library-size normalized and log-transformed. For protein data generated using 10x Visium platform and CODEX, we apply centered log-ratio (CLR) transformation to normalize expression values. For COMET protein data, background signal is first subtracted using designated background channels. We then remove background peaks by estimating the dominant background mode from the intensity distribution and clipping low-intensity values to enhance contrast. Finally, an inverse hyperbolic sine transformation is applied to stabilize variance across markers. Metabolomic data are preprocessed analogously to RNA, including library-size normalization and log transformation. After these modality-specific steps, each omics layer is standardized via Z-score normalization. Principal component analysis (PCA) is applied to reduce dimensionality, retaining the top 50 principal components as input features. In cases where a modality is measured across multiple sections, HVGs are identified by intersecting across those sections before applying the modality-specific preprocessing. To further mitigate batch effects and facilitate cross-section integration, Harmony^51^ is applied on the PCA outputs, yielding batch-corrected feature representations.

### Extraction of histology image features

Many spatial omics technologies generate paired H&E-stained histology images, which provide rich morphological context as a complementary source to molecular profiles. In addition to omics data, COSIE can incorporate H&E images as an additional modality. To effectively extract histological features, COSIE adopts the UNI model^20^, a self-supervised vision transformer built upon the ViT-Large (Patch16, 224) architecture. COSIE performs hierarchical feature extraction to capture both global and local morphological context. Specifically, for each cell, a 224 × 224-pixel image patch centered on its coordinates is extracted and fed into the UNI model. The CLS token from the final transformer layer is used as the global representation, summarizing the overall tissue characteristics around that cell. To capture local histological detail, COSIE additionally extracts the embedding corresponding to the 16 × 16-pixel central region of the patch from UNI, aligned with the cell location. The global and local representations are then concatenated to form the final image-derived feature vector for each cell. This hierarchical feature extraction ensures that both global tissue organization and fine-grained histological details are effectively captured, providing a valuable morphology-based feature for multimodal integration. The resulting image features are then reduced to 50 dimensions via PCA, consistent with the preprocessing of omics data.

Given *S* tissue sections, let *N*^*s*^ denote the number of cells in Section *s. M*^*s*^ ⊆ *M* represents the set of observed modalities of Section s, where *M* is the set of all modalities across all sections. |*M*| denotes the total number of modalities. The input data for COSIE, including all omics layers and histology-derived features, can be represented as *X* = {*X*^*s,m*^|*s* ∈ *S, m* ∈ *M*^*s*^}, where *X*^*s,m*^ represents the feature matrix of modality *m* in Section *s*.

### Cross-section linkage construction

After data preprocessing, COSIE establishes cross-section linkages to guide the subsequent integration of spatially separated sections. These linkages connect cells across different sections by leveraging both shared-modality correspondences and biologically related cross-modality relationships. Specifically, COSIE constructs two types of linkages between section pairs: (1) strong linkages between shared modalities, and (2) weak linkages across modalities. Below, we describe each type of linkages in detail.

#### Strong linkages between shared modalities

If a modality *m* is present in both Section *s*_1_ and Section*s*_2_, COSIE links cells across the two sections based on nearest neighbor (NN) matching in the feature space. Formally, for each cell in Section *s*_1_, COSIE finds its NN in Section *s*_2_ using Euclidean distance on features of modality *m*. This defines a directed linkage set from *s*_1_ to *s*_2_ for modality *m* as

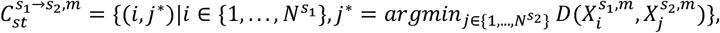

where *D*(·,·) represents the Euclidean distance measurement. To obtain the overall strong linkage from *s*_1_ to *s*_2_, we take the union over all shared modalities as

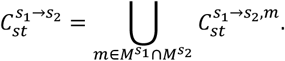

Similarly, the reciprocal linkage from *s*_2_ to *s*_1_ can be obtained as

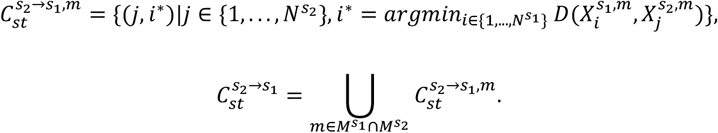

#### Weak linkages across modalities

For sections that contain different but biologically related modalities (e.g., RNA–protein, RNA–epigenomics), COSIE constructs cross-modality weak linkages leveraging the known molecular correspondences between modalities. Given a pair of Sections *s*_1_ and *s*_2_, suppose *s*_1_ contains protein data and *s*_2_ contains RNA data. We first identify shared molecular features based on known RNA-protein correspondence knowledge, resulting in two feature matrices 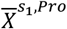 and 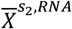. We normalize each matrix to the same total count (setting each cell’s total expression to the median across both datasets) to account for differences in scale and sequencing depth. We then apply NN search to construct weak linkages from *s*_1_ to *s*_2_ as

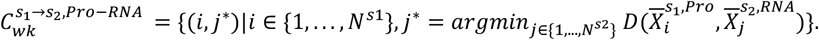

We repeat this for all relevant cross-modality pairs. Taking the union of all available weak linkages gives the overall weak linkage set from *s*_1_ to *s*_2_ as

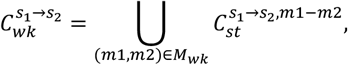

where *M*_*wk*_ denotes the set of modality pairs that have known correspondences. The reciprocal weak linkage from *s*_2_ to *s*_1_ is constructed as

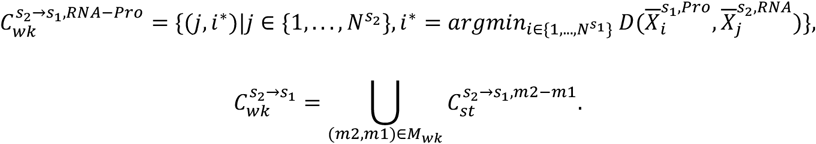

Subsequently, the complete linkages between *s*_1_ and *s*_2_ are formed by combining both strong and weak linkages as

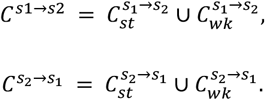

By applying this process to every pair of sections, COSIE obtains a comprehensive cross-section linkage set that captures both intra-modality correspondences and inter-modality biological relationships. The complete linkage set across all sections is defined as

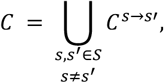

which will guide the subsequent cross-section integration process. By default, COSIE computes both strong and weak linkages for all available omics layers to ensure comprehensive cross-section alignment. However, COSIE also has the flexibility to specify custom linkage settings, allowing the exclusion of certain modality pairs that may negatively impact integration. This is particularly useful in cases where specific input modalities exhibit poor data quality, ensuring that the constructed linkages remain biologically meaningful and robust.

### Graph construction

Assuming that spatially adjacent cells and those sharing similar molecular features tend to be closely related, COSIE captures these relationships by constructing undirected cell graphs for each observed modality in each tissue section, comprising two types of edges: spatial proximity edges and feature similarity edges. To model spatial context, COSIE first constructs spatial graph based on cell coordinates. For each Section *s*, the spatial graph is defined as 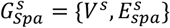, where 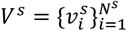 denotes the set of cell nodes. The edge set 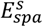 is determined by physical proximity, where an edge 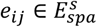 is formed if cell *j* is among the 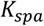 nearest neighbors of cell *i*. The collection of spatial graphs across all sections is denoted as 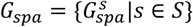.

Next, to model feature similarity, a separate graph is constructed for each available modality *m* in Section *s*, denoted as 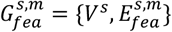. An edge 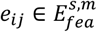 is established if cell *j* is within the *K*_*fea*_ nearest neighbors of cell *i* in the feature space. The complete set of feature graphs is denoted as 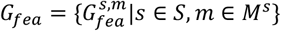.

Finally, COSIE combines both spatial and feature-based edges for each modality *m* in Section *s*, yielding a unified graph as *G*^*s,m*^ = {*V*^*s*^, *E*^*s,m*^}, where 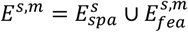. The graph structure can be represented by a binary adjacency matrix *A*^*s,m*^, where

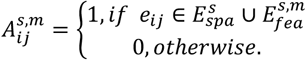

This graph construction ensures that each cell is connected to others that are either physically nearby or molecularly similar. By retaining both local spatial structure and intrinsic feature relationships, these cell graphs enable COSIE to leverage complementary information from tissue organization and multimodal profiles.

### Modality-specific encoding with graph autoencoder

With both input features and graph structures prepared, COSIE next learns a modality-specific latent representation for each cell using graph autoencoders (GAEs)^52^. Each modality *m* is assigned a dedicated GAE to aggregate information from neighboring nodes into central node. Given input feature *X*^*s,m*^ and adjacency matrix *A*^*s,m*^ for modality *m* in Section *s*, the encoding process is carried out using an *L*-layer Graph Convolutional Network (GCN)^53^. The representation at the *l*-th layer (*l* ∈ {1, …, *L*}) is given by

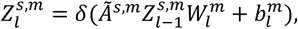

where 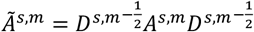 is the normalized adjacency matrix, and 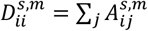 the degree matrix. 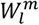 and 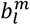 are the trainable weight matrix and bias vector for modality *m*, respectively. *δ*(·) denotes the nonlinear activation function. The input of the encoder is 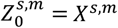, and the output 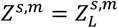 represents the modality-specific latent embedding with dimension *d*.

To reconstruct the original node features from the latent embeddings, the decoder takes *Z*^*s,m*^ as input and applies another *L*-layer GCN. Specifically, the representation at the *l*-th layer (*l* ∈ {1, …, *L*}) of decoder is computed as

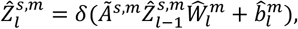

where 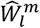 and 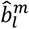 are the trainable parameters of decoder for modality *m*, and 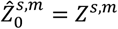. The final reconstructed feature matrix is obtained at the last layer as 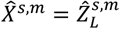.

COSIE aims to minimize the feature-wise reconstruction loss, which quantifies the difference between the reconstructed features 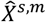 and the original input *X*^*s,m*^. The overall reconstruction loss is computed by summing across all tissue sections and observed modalities as

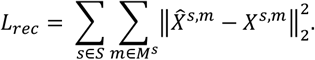

This reconstruction loss serves as the supervisory signal, guiding the encoder to learn informative modality-specific embeddings while preserving the structural relationships encoded in the graph. The complete set of learned embeddings across all observed modalities is denoted as 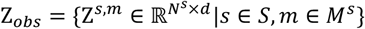.

### Information-theoretic multimodal learning within section

After obtaining the embeddings for each observed modality, COSIE then performs within-section multimodal joint learning. In this stage, modalities measured in the same section are integrated, and the representations for missing modalities are predicted. This step is essential for enhancing the representation of spatial multimodal data and inferring modalities that are not observed for a given cell. Inspired by information-theoretic multimodal learning^54^, COSIE incorporates both contrastive learning and cross-modality prediction module to improve representation consistency across different modalities.

#### Contrastive learning for paired modalities

For modalities that are jointly measured in the same section, COSIE enforces cross-modality consistency by contrastive learning. For example, consider H&E image and RNA expression present in Section *s*_1_, we have obtained their embeddings 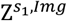 and 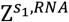 from the modality-specific encoders. The goal of contrastive learning is to maximize the mutual information between these two representations, ensuring they capture shared biological signals, while simultaneously retaining modality-specific information. This is achieved by optimizing an information-theoretic contrastive loss as

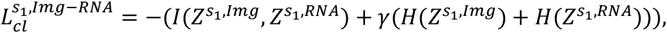

where *I*(·,·) represents the mutual information between the embeddings, and *H*(·) represents the entropy regularization term. The positive coefficient *γ* controls the balance between mutual information maximization and entropy regularization.

To approximate the mutual information, we estimate the joint probability distribution between 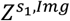 and 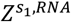, defined as

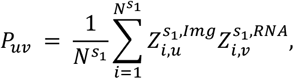

where *P* ∈ ℝ^*d*×*d*^ captures the statistical dependency between two embeddings. We then symmetrize and normalize *P* to ensure it forms a valid probability distribution as

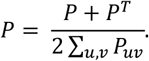

The contrastive loss is given by

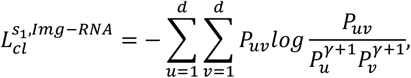

where *P*_*u*_ and *P*_*v*_ denote the marginal probability distributions, obtained by summing over the rows and columns of *P*, respectively. Since contrastive learning is applied to all paired modalities within each section, the final total contrastive loss is computed as

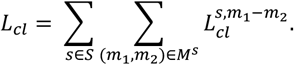

By jointly optimizing this loss, COSIE achieves robust integration between paired modalities, while preserving the intrinsic biological signals unique to each modality.

#### Cross-modality prediction via bi-directional predictors

Multimodal spatial omics data always suffer from missing modalities, where certain molecular profiles cannot be simultaneously measured across all sections. To address this issue, we introduce a bi-directional prediction module, which learns the relationship between paired modality embeddings and enables the inference of missing modality representations. Specifically, given a set of paired modalities, COSIE trains a bi-directional predictor network to model the transformations between them. For example, considering paired image and RNA embeddings in Section *s*_1_ again, we define the bi-directional predictors between image and RNA as

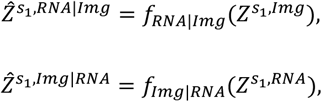

where *f*_*RNA*|*Img*_(·) and *f*_*Img*|*RNA*_(·) represent the predictors that transform embeddings from image to RNA and RNA to image, respectively. We implement these predictors by multilayer perceptron (MLP), which take the embedding from one modality as input and generate a transformed representation in the target modality space 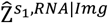 denotes the predicted RNA embedding, and 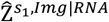 denotes the predicted image embedding. COSIE trains these predictors by minimizing the cross-modality prediction loss that penalizes the difference between the predicted embeddings and the actual embeddings. For the image-RNA example in Section *s*_1_, the prediction loss is formulated as

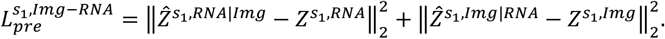

COSIE extends this loss across all sections and all pairs of modalities that are considered predictable from one another. The total prediction loss is given by

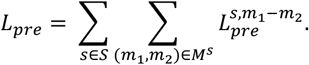

This prediction module provides two key advantages. First, the prediction loss can be interpreted from an information-theoretic perspective. The bi-directional prediction process is intrinsically to minimize the conditional entropy 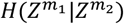 and 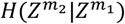 between two paired modalities *m*_1_ and *m*_2_. By learning predictors that accurately reconstruct one modality from the other, COSIE encourages the representation 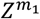 to be fully determined by 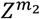 and vice versa, thereby reducing 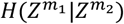 and 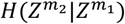. This complements the contrastive loss, which maximizes the mutual information 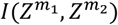. These two objectives benefit each other and together lead to a more accurate cell representation.

Second, the trained predictors enable the prediction of missing modality embedding, ensuring a complete multimodal representation for each cell. For each missing modality, its predicted embedding is inferred from available modality embeddings using the trained predictors within the same section. If multiple observed modalities can predict the missing one, COSIE uses all relevant predictors and averages the resulting predictions to obtain the predicted embedding. If no predictors are available for a missing modality, an all-zero matrix is assigned. We denote the set of predicted embeddings as 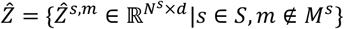.

Finally, for each cell we concatenate its learned encoder embedding and predicted embeddings for missing modalities as

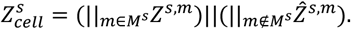

The set of cell embeddings across all sections is then represented as 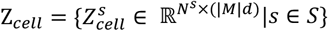.Here || denotes the concatenation operation along the feature dimension. This formulation ensures that every cell retains a complete multimodal representation, allowing for a unified and comparable embedding space across different sections, even when certain modalities are missing.

### Cell neighborhood-level triplet learning across sections

To align cells across different tissue sections in a universal feature space, COSIE subsequently performs cross-section integration based on the previously constructed cross-section linkage. This process is conducted at the cell neighborhood level, ensuring that local spatial structures are well preserved. Specifically, for Section *s*, we leverage the pre-established spatial edges 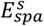 to derive cell neighborhood embeddings. Given a cell *i* in Section *s*, its cell neighborhood embedding is computed as

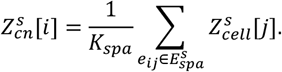

The complete set of cell neighborhood embeddings across all sections is then defined as 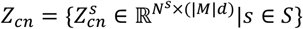.

By incorporating cell neighborhood embeddings, COSIE enables cells with similar spatial contexts to exhibit consistent representations across sections, facilitating robust cross-section alignment. Next, we propose a triplet loss that leverages both the previously constructed cross-section linkages and cell neighborhood embeddings. For a given pair of Sections *s*_1_ and *s*_2_, and taking cells in *s*_1_ as anchors, the positive samples are cells in *s*_2_ that are linked to the anchor in cross-section linkage 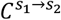, while the negative cells are randomly selected from Section *s*_1_. The triplet loss can be formulated as

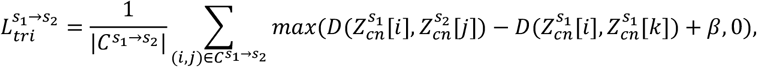

where 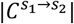 is the number of cross-section linkages in 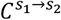 and *k* denotes the randomly sampled negative cell from Section *s*_1_. *β* is the margin hyperparameter that ensures the anchor-positive pairs remain closer than the anchor-negative pairs.

Similarly, when treating cells in *s*_2_ as anchors and linking them to corresponding positive cells in *s*_1_ through 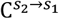, the triplet loss is computed symmetrically. By extending across all section pairs, the final triplet loss is given by

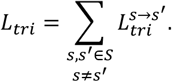

This loss encourages the encoder to produce biologically meaningful and batch-corrected embeddings by pulling linked cell pairs closer and pushing unrelated pairs apart. At the same time, local spatial relationship is well-preserved through the use of neighborhood embeddings. After training, cells with analogous phenotypes in different sections will end up close in the unified embedding space, whereas dissimilar cells remain apart.

### Overall loss function

Finally, the overall objection function of COSIE integrates all the components and is expressed as

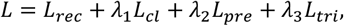

where *λ*_1_, *λ*_2_ and *λ*_3_ are the balanced factors. The training process of COSIE follows a staged strategy to stabilize training as new objectives are gradually introduced: Initially, both *λ*_2_ and *λ*_3_ are set to 0, and only the graph autoencoder module is trained. At this stage, the model focuses on learning meaningful latent cell embeddings from raw input data while enforcing within-section consistency through contrastive learning. Once the embeddings get more stable (we set epoch 100), the cross-modality prediction module is introduced by setting *λ*_2_ to 0.2. This enables COSIE to infer missing embeddings across different modalities based on learned relationships. From epoch 200, as the prediction module becomes stable, we incorporate the triplet loss by setting *λ*_3_ to 1.0. At this stage, missing cell representations are completed, and neighborhood-level triplet learning is leveraged to integrate multiple sections while maintaining local spatial coherence. After training, the final representation of each cell Z_*final*_ is obtained by averaging its cell embedding and cell-neighborhood embedding, ensuring the preservation of both intrinsic biological information and spatial context within its microenvironment. This unified representation is utilized for various downstream analyses.

### Missing omics data prediction

With the unified embedding space well established, COSIE enables the prediction of missing omics data by leveraging cross-section cell similarities. Specifically, for a given section with missing modality *m*, COSIE performs KNN search in the embedding space to retrieve *K*_*imp*_ cells from other sections that have observed data for modality *m*. The missing data is then predicted by averaging the expression values of the retrieved neighbors. This embedding-based retrieval strategy enables COSIE to borrow information from highly similar cells, regardless of their spatial origin, and flexibly complete partially observed modalities.

### Scaling to large-scale datasets

Recent advances in spatial omics technologies have led to increasingly large datasets with hundreds of thousands or even millions of cells, posing significant computational challenges for graph-based learning frameworks. To ensure scalability, COSIE adopts a spatially aware subgraph partitioning and batch-wise training strategy.

Specifically, each tissue section is divided into several spatial subregions along the x and y axes. All modalities within a section are partitioned in a consistent manner, ensuring that each subgraph contains matched data. COSIE then constructs spatial and feature similarity graphs independently within each subgraph. Instead of computing linkages between entire sections, COSIE now computes cross-subgraph linkages between all pairs of subgraphs across different sections. During training, COSIE performs optimization in a batch-wise fashion. This subgraph-level batching drastically reduces memory usage while maintaining learning effectiveness, allowing COSIE to process large-scale datasets. In the final evaluation stage, embeddings are computed on the full graph for each section to ensure global consistency. To further alleviate data sparsity and reduce computational complexity, COSIE supports spatial meta-cell construction by aggregating every four spatially adjacent cells into a single meta-cell. This strategy accelerates training on large-scale datasets while preserving local spatial architecture.

### Hyperparameters

COSIE adopts a consistent set of hyperparameters across all datasets. For the graph construction, we use *K*_*spa*_ = 5 for the spatial graph, and *K*_*fea*_ = 30 for the feature similarity graph. COSIE is trained for 600 epochs using the Adam optimizer with a learning rate of 0.0001. The coefficient γ in the contrastive loss is set to 5. The balanced factors in the overall loss function are set to *λ*_1_ = 0.1, *λ*_2_ = 0.2, and *λ*_3_ = 1.0. The margin hyperparameter *β* is set to 1. The graph autoencoder consists of two layers with hidden dimensions [256, 128]. The bi-directional predictors are all implemented with MLP with hidden dimensions of [512, 512].

### Clustering analysis

All clustering was performed using the K-means algorithm. To examine the relationships among clusters, we computed the centroid of each cluster by averaging the representations of all cells assigned to that cluster. These centroids reflect the global embedding characteristics of each cluster within the integrated space. Hierarchical clustering was then performed on the centroid matrix using average linkage via the *scipy*.*cluster*.*hierarchy*.*linkage* function. The resulting dendrogram, generated using *scipy*.*cluster*.*hierarchy*.*dendrogram*, was used to visualize the structural similarity between clusters. Clusters with closely positioned centroids were interpreted as having similar spatial characteristics. This dendrogram guided post hoc cluster merging, in which clusters with closely located centroids were aggregated.

### Spatial registration across modalities

To perform spatial alignment between H&E images and other spatial omics data, we employed affine transformation based on manually annotated landmark pairs. The affine transformation matrix was estimated using the *cv2*.*findHomography* function with the RANSAC algorithm to ensure robustness against outliers. The resulting transformation was then applied to the source coordinates using *cv2*.*perspectiveTransform*, enabling all modalities to be spatially mapped into a common coordinate system.

### Differential expression and functional enrichment analysis

We performed differential gene expression analysis using the Wilcoxon rank-sum test as implemented in the *scanpy*.*tl*.*rank_genes_groups* function^55^. Cells were grouped based on their cluster labels. Functional enrichment analysis based on detected differentially expressed genes was carried out using g:Profiler^56^.

### Pseudotime analysis

Trajectory inference on the mouse embryo ATAC data was performed using Slingshot^57^ (v2.16.01), based on principal component analysis (PCA). For Section 1 and Section 2, we used predicted and observed ATAC data, respectively. For the lung cancer dataset, we employed diffusion pseudotime (DPT)^58^ analysis via Scanpy to accommodate the large data scale. After dimensionality reduction using PCA, we constructed a k-nearest neighbor graph (k=15, 30 PCs) and applied Leiden clustering (resolution=0.5). DPT was then computed using 10 diffusion components, with the root defined as the normal lung cluster.

### Copy number variation inference

We used the SpatialInferCNV^46^ R package to infer genome-wide copy number variations (CNVs) from Visium spatial transcriptomics data for the LUAD sample, following the official tutorial (https://aerickso.github.io/SpatialInferCNV/). Clones were identified based on clustering of CNV profiles, and a subset of spots with minimal CNV signal and consistent with histologically normal epithelium was selected as the reference. Final CNV inference was then performed using this reference population. To reconstruct clonal phylogenies, we computed pairwise distances between clone-level consensus CNV profiles and applied the Neighbor-Joining method^59^. Clone identities were subsequently assigned to all spots based on their positions within the resulting dendrogram.

### Survival analysis

Survival analysis was conducted using the interactive web-based platform GEPIA2^60^ (http://gepia2.cancer-pku.cn/#survival), based on the TCGA LUAD cohort (**Fig. 6**, human lung cancer-related). Kaplan–Meier curves were generated to compare overall survival between patients with high versus low gene expression. A 75% percentile cutoff was applied to stratify patients into high and low expression groups. Statistical significance was assessed using the log-rank test. The dotted lines in the survival plots indicate the 95% confidence intervals.

### Competing methods

StabMap^12^ was implemented following the tutorial (https://marionilab.github.io/StabMap/articles/stabMap_PBMC_Multiome.html). After obtaining the StabMap embeddings, we further applied Harmony for batch correction, as recommended by the authors. scMoMaT^13^ was implemented using its Python package (v0.2.2), following the tutorial provided at GitHub (https://github.com/PeterZZQ/scMoMaT/blob/main/demo_scmomat.ipynb). SpaMosaic^11^ was implemented according to its documentation (https://spamosaic.readthedocs.io/en/latest/). StabMap used the same input preprocessing as COSIE, while scMoMaT and SpaMosaic employed their own custom normalization. All methods produced low-dimensional embeddings, which were subsequently used for clustering and visualization.

### Training and running time

All COSIE experiments were conducted using a single NVIDIA A100 GPU with 80 GB memory. Model training time of COSIE and competing methods on each dataset was summarized in **Supplementary Fig. 20**. Among all datasets, the four-section integration in human lung cancer dataset was the largest, consisting of over 3 million superpixels across tissue sections. COSIE completed the training step on this dataset within 23 minutes, with a peak GPU memory usage of approximately 65 GB.

### Periodontal disease and diffuse large B-cell lymphoma data generation

Periodontal disease and diffuse large B-cell lymphoma (DLBCL) datasets were generated following the IN-DEPTH strategy^7^. For the periodontal disease data, formalin-fixed paraffin-embedded (FFPE) tissues were sectioned at 5 μm thickness on SuperFrost 697 glass slides (VWR, 48311-703) and were provided by D.M.K. from Harvard Dental School (IRB# 22-0587). Two adjacent sections were prepared: Section 1 included paired H&E staining and Visium HD profiling, while Section 2 included paired CODEX and Visium HD data. For all protein markers in the periodontal dataset, images were derived from the 8-bit final QPTIFF files generated by the PhenoCycler Fusion software, with default blank cycle subtraction applied.

For the DLBCL dataset, FFPE tissues were also sectioned at 5 μm thickness on SuperFrost 697 glass slides. Two samples were analyzed in this study, each with H&E and CODEX imaging at the full core level, and CosMx profiling at the field-of-view (FOV) level. The CosMx data were uploaded to the AtoMx platform, from which decoded transcripts and their corresponding spatial coordinates were exported. To enable spatial alignment across imaging modalities, VALIS^61^ was used to register CosMx transcriptomic data and H&E images to the spatial coordinate system of the CODEX proteomic dataset, ensuring consistent spatial referencing across all modalities.

### Atherosclerosis Xenium and CODEX data generation

#### Xenium sample preparation

Spatial single-cell transcriptomic profiling of human atherosclerosis tissue was performed using the Xenium *In Situ* platform (10x Genomics). FFPE tissue blocks were sectioned at 5 μm and mounted onto Xenium slides. Sample processing was carried out according to the manufacturer’s protocol, including deparaffinization, decrosslinking, probe hybridization, ligation, and signal amplification over a three-day workflow. Xenium assays were run using instrument software version 1.5.1.2 or 1.7.6.0 and processed using analysis versions 1.5.0.3 or 1.7.1.0. After transcriptomic imaging, H&E staining was performed following the recommended post-Xenium protocol. Two immediately adjacent sections were profiled using the Xenium platform, each employing a fully customized gene panel. Genes were selected based on their relevance to key cellular functions, disease-associated pathways, and differentially expressed genes identified from prior datasets. Panel 1 included 314 genes and Panel 2 included 345 genes, with 111 genes shared between them to capture core cellular processes.

#### CODEX sample preparation

CODEX section is approximately 35 µm away from the Xenium sections. FFPE tissue sections were mounted on coverslips and processed using the PhenoCycler™-Fusion platform (Akoya Biosciences) according to the manufacturer’s protocols. Sections were deparaffinized, rehydrated, and subjected to heat-induced epitope retrieval using Akoya’s pH 9 antigen retrieval buffer. PhenoCycler experiments were designed using the PhenoCycler Experiment designer 2.1.0 (Akoya Biosciences). Tissues were stained with the DNA-barcoded antibody panel (42 proteins in total). After staining, mounted chambers were loaded onto the PhenoImager® Fusion system. Fluorescently labeled oligonucleotide reporters were hybridized in sequential cycles to visualize the barcoded antibodies, and signal acquisition was performed for each round. Raw image data were processed using the software Fusion 2.3.1 (Akoya Biosciences), including image alignment, background subtraction, and signal decoding.

### Mouse coronal brain spatial MALDI-Visium data generation

The mouse coronal brain dataset includes two sections. Section 1 was generated in MISO^32^, and section 2 was generated following the same experimental procedure. Visium slides (10x Genomics) were physically modified to be compatible with the MALDI imaging mass spectrometry platform by trimming and sanding the edge, allowing them to fit into the Bruker Daltonics slide adaptor. Fresh-frozen, non-perfused brain tissues from 9-month-old mice were cryosectioned at 12 µm thickness and mounted onto the adapted slides. N-(1-naphthyl) ethylenediamine dihydrochloride (NEDC) was used as the MALDI matrix at a concentration of 10 mg/ml in 70:30 (v:v) methanol:water and sprayed on the tissue sections using an HTX-TM sprayer (HTX Technologies) as described previously^32^. Mass spectrometry imaging was performed as described in detail previously^32^. Briefly, prepared tissue slices on Visium slides were imaged using a MALDI-2 timsTOF fleX instrument (Bruker Daltonics) with 20 µm spatial resolution, operating in negative ion mode over an m/z range of 100–1000 and without engaging trapped ion mobility separation (TIMS). Spatial transcriptomics data were generated using the 10x Genomics Visium platform and Space Ranger pipeline (10x Genomics) was utilized to process the raw sequencing data. H&E staining was performed for histological imaging using a Leica Aperio slide scanner.

### Gastric cancer Visium and COMET data generation

#### Spatial Transcriptomics data generation using Visium

Spatial transcriptomics was performed on FFPE gastric tissue sections using the 10x Genomics Visium CytAssist platform, following the manufacturer’s protocols. Archived H&E-stained slides were reviewed by a pathologist to assess tissue quality, including preservation, orientation, and the absence of folds or contaminants. Based on histological evaluation, a single region of interest (ROI) per sample, measuring 11 × 11 mm, was selected according to tumor content, tissue integrity, and anatomical relevance. Five-micrometer-thick sections from each FFPE block were mounted on Superfrost Plus slides, stained with H&E using a Leica Autostainer, and scanned at high resolution with the Aperio AT2 system. The Visium CytAssist instrument was then used to transfer the transcriptomic capture area onto the Visium Gene Expression slide via precise placement of the gasket over the selected ROI. After tissue permeabilization, targets were released and hybridized to barcode oligoprobes with defined XY coordinates on the slide, yielding over 14,000 barcodes per section. Library preparation was carried out using the Visium CytAssist Spatial Gene Expression FFPE Library Construction Kit with the Transcriptome Probe Set. The resulting libraries were quality-checked and sequenced on an Illumina NovaSeq 6000 platform. Pathological annotation was conducted manually in Loupe Browser (10x Genomics, v8.0.0) based on the high-resolution H&E images.

#### Automated seqIF-based spatial proteomics and imaging on COMET

Sequential immunofluorescence (seqIF) was conducted using the COMET platform (Lunaphore Technologies). FFPE tissue sections were processed through baking, deparaffinization, and rehydration, followed by autofluorescence quenching and antigen retrieval (pH9 EZ AR 2 Elegance, Biogenex), in accordance with optimized in-house protocols. Slides were subsequently loaded onto the COMET Stainer and processed per the manufacturer’s guidelines. Samples were subjected to iterative cycles of immunostaining, involving incubation with primary and secondary antibodies, imaging, and antibody elution. Each staining round was followed by image acquisition before proceeding to the next cycle, as previously described^62,63^. The antibody panels (19 proteins) were customized for the gastric cancer tissue. Primary antibody incubations were carried out for 8 minutes per round. Upon completion of the staining cycles, multi-layer ome.tiff images from each round were automatically stitched and aligned. Background fluorescence was computationally subtracted, and the images were visualized and analyzed using Lunaphore Viewer.

### Lung cancer 10x Visium spatial gene and protein data generation

Spatial gene and protein data were generated from four surgically resected human LUAD samples (P11, P12, P15, and P24) using the 10x Genomics Visium spatial gene and protein platform. For sample P11, single-cell spatial transcriptomic profile was obtained from an adjacent Xenium section. These data enabled multimodal spatial characterization of LUAD lesions, including both invasive components and surrounding histological structures.

#### Visium CytAssist processing and H&E imaging

FFPE tissue blocks from each LUAD case (P11, P12, P15, P24) were sectioned at 5 μm thickness and mounted on standard glass slides. H&E staining was performed using the Leica ST5010 Autostainer XL according to manufacturer protocols. H&E slides were scanned at 40× magnification using an Aperio ScanScope Turbo (Leica Microsystems) to produce high-resolution whole-slide images. Prior to Visium CytAssist spatial profiling, each slide was carefully reviewed by a board-certified pathologist to confirm tissue quality and the presence of tumor and peritumoral regions.

Spatial transcriptome and protein (35 proteins) profiling were performed using the Visium CytAssist Gene and Protein Expression assay (10x Genomics), following manufacturer-recommended protocols. On average, ∼11,000 barcoded capture spots were obtained per sample. Libraries were sequenced on an Illumina NovaSeq platform to a target depth of approximately 65,000 reads per spot. Raw sequencing data were processed using Space Ranger (v2.0.0) and aligned to the GRCh38 human reference genome. Spots with low quality were excluded. Manual spot annotation based on H&E morphology was performed using Loupe Browser (v7.1), and regions annotated as “Space”, “Tissue artifact”, or “Foreign body” were filtered out. To enable single-cell resolution analysis, we applied iStar to enhance both gene and protein data into super-resolution (16 × 16 pixels) across the tissue sections.

#### Xenium data generation

To supplement the Visium-based profiling, an adjacent 5 μm FFPE tissue section from sample P11 was analyzed using the Xenium *In Situ* platform (10x Genomics) to achieve spatially resolved single-cell transcriptomic profiling. The tissue section was processed and hybridized according to the manufacturer’s protocol. Xenium imaging and molecular decoding were performed using Xenium Analyzer, resulting in single-cell resolution RNA quantification for a predefined gene panel with 5101 genes.

## Acknowledgments

M.L. was partly supported by the National Institutes of Health (NIH) grants R01HG013185, R01LM014592, R01HL171595, U01CA294518, and U19NS135528. Y.D. was partly supported by the Packard Fellowship for Science and Engineering and NIH grant DP2AI177913. S.J. was supported in part by NIH grants DP2AI171139, P01AI177687, R01AI149672, R01GM152585, U24CA224331, R01NS139479, a Gilead’s Research Scholars Program in Hematologic Malignancies, the Broad Next Generation Award, the Dye Family Foundation, and the Bridge Project, a partnership between the Koch Institute for Integrative Cancer Research at MIT and the Dana-Farber/Harvard Cancer Center.

## Author contributions

This study was conceived of and led by M.L. W.L. designed the model and algorithm, implemented the COSIE software and led data analyses with input from M.L. W.L. and L.M. conducted benchmarking analyses. L.M. processed the epigenomics data, carried out pseudo time analyses and contributed to the COSIE documentation. L.W. and Y.L. generated and processed the gastric cancer data. H.K. and F.P. generated and processed the lung cancer data. L.W., H.K., Y.L. and F.P. provided input for cancer data analysis. L.Maegdefessel and N.S. generated, processed and annotated the atherosclerosis data. S.J., A.K.S, D.M.K., W.W., S.P.T.Y., A.B., Y.W.C. and C.Y.C generated and processed the periodontal and B cell lymphoma data. W.W. carried out the multimodal registration for the B cell lymphoma data. S.P.T.Y. conducted CODEX-based phenotyping for the periodontal data. H.Y. performed CODEX image preprocessing for the atherosclerosis data. A.S. and X.Y. contributed to H&E preprocessing and feature extraction. K.T. polished the figures. S.J. developed an approach to align clusters obtained from different methods. S.J. and Z.C. provided input on data analysis and presentation. M.L.L. processed the spatial MALDI-Visium data in mouse brain. L.G. annotated the COSIE-identified macrophages in the lung cancer data. I.L. and L.M.S.S. annotated the gastric cancer data. M.P.R. provided input for the atherosclerosis data analysis. X.Q. annotated the mouse hippocampus region and provided input for the spatial MALDI-Visium data analysis. C.A.T., N.B., L.Z.S., and J.D.R. generated the mouse hippocampus spatial MALDI-Visium data. Y.D. and L.M. provided input for the Spatial-Mux-seq data analysis. W.L. and M.L. wrote the paper with feedback from the other co-authors.

## Competing financial interests

M.L. receives research funding from Biogen Inc. unrelated to the current manuscript. M.L. is a co-founder of OmicPath AI LLC. Y.D. is the scientific advisor of AtlasXomics Inc. S.J. is a co-founder of Elucidate Bio Inc, and has received research support from Roche and Novartis unrelated to the current manuscript. L.M.S.S. reports research support from Theolytics, advisory role/consulting fees from BioNTech, and travel support from 10x Genomics, all outside the scope of this work.

